# A novel post-translational proteomics platform identifies neurite outgrowth impairments in Parkinson’s disease *GBA-N370S* dopamine neurons

**DOI:** 10.1101/2021.06.30.450333

**Authors:** Helle Bogetofte, Brent J. Ryan, Pia Jensen, Dana L.E. Vergoossen, Mike B. Barnkob, Lisa Kiani, Uroosa Chughtai, Janine Brandes, Jane Vowles, Fiona Bunn, Peter Kilfeather, Hugo J.R. Fernandes, Tara Caffrey, Morten Meyer, Sally A. Cowley, Martin R. Larsen, Richard Wade-Martins

## Abstract

The causes of Parkinson’s disease (PD) likely involve complex interactions between environmental factors and susceptibility genes with variants at the *GBA* locus encoding the glucocerebrosidase (GCase) enzyme being the strongest common genetic risk factor for PD. To understand *GBA*-related disease mechanisms, we used a novel multipart-enrichment proteomics and post-translational modification workflow to simultaneously identify peptides with phosphorylation, reversible cysteine-modifications or sialylated N-linked glycosylation, alongside unmodified proteins.

We identified large numbers of dysregulated proteins and post-translational modifications (PTMs) in heterozygous *GBA*-*N370S* PD patient induced pluripotent stem cells (iPSC)-derived dopamine neurons. Alterations in glycosylation status of lysosomal proteins identified disturbances in the autophagy-lysosomal pathway, concurrent with upstream perturbations in mTOR phosphorylation and activity in *GBA-N370S* iPSC-dopamine neurons. In addition, the strategy revealed several native and modified proteins encoded by PD-associated genes to be dysregulated in *GBA-N370S* neurons, enhancing our understanding of the wider role of *GBA* mutations on the neuronal proteome. Integrated pathway analysis of all datasets revealed impaired neuritogenesis in *GBA-N370S* PD iPSC-dopamine neurons and identified tau (*MAPT*) as a key mediator of this process. Using a functional assay, we confirmed neurite outgrowth deficits in *GBA-N370S* PD neurons and a central role for tau in this process. Furthermore, pharmacological restoration of GCase activity in *GBA-N370S* PD patient neurons rescued the neurite outgrowth deficit. Overall, this study demonstrates the potential of PTMomics to elucidate novel neurodegeneration-associated pathways and identify phenotypes and potential drug targets in complex disease models.

## Introduction

Heterozygous mutations in the glucocerebrosidase gene (*GBA)* are the strongest common genetic risk factors for Parkinson’s disease (PD) present in around 5-10% of PD patients (2, 3), resulting in lower age of onset and exacerbating disease progression, including an increased risk of dementia (4, 5). However, the exact mechanisms leading from dysfunction of the enzyme glucocerebrosidase (GCase) encoded by *GBA* to PD pathogenesis and neurodegeneration remain unclear. GCase is a lysosomal enzyme which degrades glucosylceramide into glucose and ceramide. Homozygous mutations in *GBA* lead to severe GCase deficiency and glucosylceramide accumulation in lysosomes preventing their normal function (6) causing the lysosomal storage disorder Gaucher’s disease (GD). Heterozygous *GBA* mutation carriers have a 10-30% risk of developing PD (7, 8). Over 300 different disease-associated changes in *GBA* have been identified, of which the N370S mutation is the most common (9).

Studies of how *GBA* mutations may lead to PD pathogenesis have generated diverse hypotheses involving either loss-of-function or toxic gain-of-function mechanisms of a number of cellular pathways including the autophagy-lysosome pathway (ALP), ER stress or lipid dyshomeostasis (10, 11). Decreased GCase activity has been demonstrated in patients with genetic and sporadic PD, and also as a consequence of aging, suggesting that loss of enzyme activity may be causal for disease (12, 13). Human cell and animal models of GCase deficiency, obtained by genetic knockout or chemical inhibition, highlighted ALP dysfunction and accumulation of α-synuclein (α-syn) (14–16). This could be the result of reduced ALP function hindering the degradation of α-syn either directly due to a loss of GCase activity or due to a toxic gain-of-function of mutant GCase (11, 17–20).

Induced pluripotent stem cell (iPSC)-derived dopamine neurons from PD patients have been fundamental in understanding the molecular pathology of PD. Studies using iPSCs derived from patients with *GBA* mutations show increased dopamine oxidation, decreased lysosomal function, accumulation of glycosphingolipids including glucosylceramide, and build-up of α-syn (17–19). Furthermore, we have previously demonstrated *GBA*-*N370S* phenotypes including ALP dysfunction, augmented ER stress response and increased α-syn release in patient iPSC-derived dopamine neurons (11, 20).

Proteomic and single-cell transcriptomic strategies have been previously used to characterise dysfunction in iPSC-derived neurons from patients with *GBA* mutations, highlighting the utility of ‘omics approaches (18, 20). However, in order to fully understand the cellular effects of *GBA* mutations an unbiased and global characterisation of the neuronal proteome is required. Furthermore, given a large proportion of regulation of protein activity, localisation and function is not controlled by changes in protein levels, but through post-translational modifications (PTMs) (21), simultaneous assessment of multiple protein PTMs is essential in understanding complex diseases mechanisms and models.

Numerous PTMs have been associated with cellular dysfunction and PD pathology. Phosphorylation, which is the most studied PTM, has been demonstrated to be involved in regulating activity of major PD-related pathways such as ALP and mitophagy (22). In addition, the phosphorylation states of key proteins involved in neurodegeneration such as α-syn and tau can affect their propensity for aggregation (23). Another major contributor to PD pathology is oxidative stress as increased levels of reactive oxidative/nitrosative species cause reversible and irreversible modifications of redox-sensitive cysteine residues, affecting both the structure and function of proteins (24, 25). Additionally, N-linked glycosylation of proteins is a highly relevant PTM for understanding the proteome, with a large number of proteins shown to have N-linked glycosylation (26). The prevalence of this modification and the role sialylated N-linked glycosylation plays in neural development and function, including in processing and transport of GCase and other key lysosomal proteins through the ER and Golgi (3, 27, 28), highlights the need to assess the abundance of sialylated N-linked glycosylation in the proteome of disease models. We have previously examined the effect of knockdown of the PD-linked gene *PRKN (PARK2)* on the proteome of iPSC-derived neurons focusing on phosphorylation and oxidised cysteine-containing proteins, resulting in identification of a number of phenotypes (29–31).

Applying a novel multi-part enrichment strategy (TiCPG) (1), we have isolated and quantified peptides with phosphorylation, reversible cysteine-modification or sialylated N-linked glycosylation, simultaneously. We applied this methodology to iPSC-derived dopamine neurons from *GBA*-*N370S* PD patients and healthy age-matched controls, identifying large numbers of dysregulated native and modified proteins. Quantification of numerous sialylated glycopeptides, from lysosomal proteins identified disturbances in the ALP which coincided with upstream perturbations in mTOR levels. Integrated pathway analysis of all PTMomic datasets revealed an enrichment of dysregulated native/PTM proteins related to neuritogenesis, which we validated in functional assays, showing significant defects in neurite outgrowth, a novel phenotype in *GBA-N370S* patient neurons. Knockdown of tau, a predicted upstream regulator of *GBA*-induced dysfunction, caused a significant decrease in neurite outgrowth, whereas normalising GCase activity using a small molecule chaperone significantly rescued the neurite outgrowth defects. Together, this study highlights the potential of this novel workflow for identifying phenotypes and their underlying biology as well as potential drug targets in disease models.

## Results

### Proteomic analysis reveals the post-translational proteome of human midbrain dopaminergic neurons

To assess the proteome and PTMome in human dopaminergic neuronal cultures, iPSC lines from four age and sex matched *GBA-N370S* mutation carriers diagnosed with PD and four healthy control iPSC lines were differentiated into dopaminergic neuronal cultures with comparable efficiency (Fig. 1A, Fig. S1 and Table S1). Neurons derived from *GBA* mutation carriers exhibited a 50% decrease in GCase activity (Fig. 1B).

**Fig 1:**
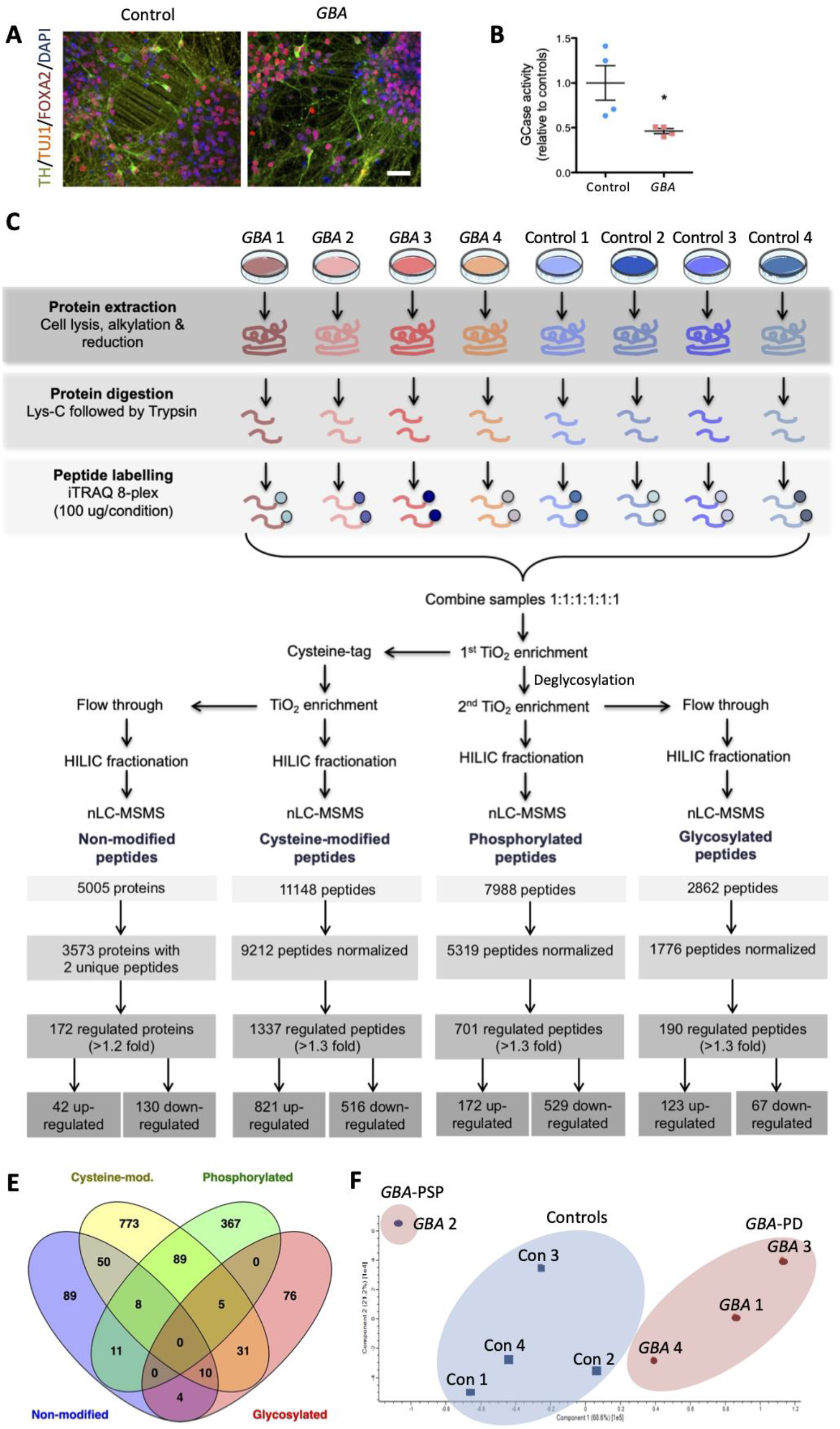
Proteomic analysis on *GBA* and control iPSC-dopamine neurons achieves clear patient stratification. A) Immunofluorescence staining for tyrosine hydroxylase (TH, green), β-III-tubulin (TUJ1, yellow), and FOXA2 (red) on differentiation day 35 *GBA* patient and control neurons. Nuclei stained with DAPI (blue). Scale bar = 50 μm. B) Glucocerebrosidase (GCase) enzyme activity, relative to average of controls, of the *GBA* patient and control iPSC-dopamine neurons included in the proteomic analysis (n = 4 *GBA* patients and 4 controls; Mean ± SEM. *P≤0.05 (Student’s *t*-test). C) Schematic representation of the preparation and enrichment workflow for the proteomic analysis of *GBA* patient and control iPSC-dopamine neuron cell lysates. D) Overview of the identified non-modified proteins and phosphorylated/ glycosylated/cysteine-modified peptides. Normalised peptides represent number of individual phosphorylated/glycosylated/cysteine-modified peptides normalized to levels of the corresponding non-modified proteins. E) Venn diagram showing the overlap between the resulting regulated proteins and modified peptides in the four groups. F) Principal component analysis (PCA) plot based on the protein expression data from iPSC-dopamine neurons derived from the four *GBA* patients (circles, *GBA* 2; *GBA*-PSP in purple) and healthy controls (squares).

Dopaminergic neuronal cultures were harvested and peptides derived by tryptic digestion from neural proteins were labelled to allow multiplexing. The peptide mixture was subsequently subjected to the sequential enrichment strategy TICPG, followed by pre-fractionation and subsequent LC-MS/MS (Fig. 1C). The mass spectrometry analysis successfully isolated, identified and quantified more than 5,000 proteins with high confidence across all 8 samples. In addition, we obtained the first ever global characterisation of several important subtypes of PTMs in human midbrain neurons. In total, 7,988 phosphorylation sites on 3,092 proteins and 2,862 sialylated N-linked glycosites on 1,055 proteins were identified. In addition, 11,148 reversible cysteine-modifications on 4,456 proteins were detected (Fig. 1D), providing a comprehensive reference dataset of the proteome and PTMome in human iPSC-dopamine neurons.

### Proteomic signatures of *GBA*-PD iPSC-dopamine neurons are specific and distinct from healthy controls

Analysis of the proteome resulted in the identification of 172 proteins that were significantly differentially expressed in *GBA-N370S* neurons with a >1.2-fold change in abundance compared to controls (Fig. 1D, Table S2A-B). To understand the contribution of PTMs to cellular dysfunction in *GBA-N370S* mutation carriers, levels of PTM peptides were normalised to the level of the corresponding non-modified protein. This analysis resulted in more than 20,000 normalised PTMs (Table S2). Of these, levels of 1,337 reversible cysteine-modifications, 701 phosphorylation sites and 190 sialylated glycosites were altered by more than 1.3-fold in *GBA-N370S* iPSC-dopamine neurons and these PTMs were included in downstream analyses (Fig 1D, Table S2C-H).

Comparing the proteins which were differentially expressed and/or showed changes in PTM levels, a total of 201 proteins were found to be regulated in at least two of the datasets, signifying increased regulation associated with *GBA* mutation-related disease mechanisms (Fig. 1E).

Principal component analysis (PCA) of all non-modified proteins separated the four controls from three *GBA* iPSC-dopamine neurons, with neurons from the *GBA* patient 2 line clearly separating from the others (Fig. 1F). This was confirmed by hierarchical clustering of the 500 most abundant proteins (Fig. S2A). Similarly, PCA and hierarchical clustering of the post-translationally modified proteins separated patients from control iPSC-dopamine neurons and segregated *GBA* patient 2 from the remaining *GBA* neurons (Fig. S3). As previously described elsewhere (23), post-hoc clinical follow-up of *GBA* patient 2 revealed poor L-DOPA responsiveness and altered disease course with the individual later diagnosed with progressive supranuclear palsy (PSP), and not Parkinson’s disease (20).

Comparing protein expression between the *GBA* patient 2 iPSC-dopamine neurons and the four healthy controls revealed levels of 250 proteins to be significantly altered (Table S3). Pathway analysis of these proteins showed an enrichment of proteins involved in extracellular matrix organisation and neurotransmitter release cycle, in addition to regulation of cell biology by calpain proteases, which are known regulators of tau phosphorylation and fragmentation (Fig. S2B-C) (35, 36). Given the differential diagnosis, *GBA* patient 2 was excluded from analysis of the remaining *GBA* mutation carriers all of whom had a confirmed PD diagnosis and termed *GBA*-PD.

These data demonstrate the ability of our comprehensive analysis to differentiate between iPSC-dopamine neurons derived from *GBA-N370S* carriers and controls and from patients with different forms of neurodegenerative diseases.

### Assessment of sialylated glycopeptides reveals widespread changes in the lysosomal proteome of *GBA-*PD iPSC-dopamine neurons

Glycosylation is an abundant PTM, particularly on membrane bound and lysosomal proteins, suggesting their levels may be relevant in *GBA*-PD (11, 14, 37). Hierarchical clustering and pathway analysis of the complete sialylated glycoprotein dataset demonstrated coverage of proteins involved in cell motility, neurite outgrowth and lysosomal processes, and PCA analysis of these peptides revealed modest separation of *GBA*-PD and control neurons, in addition to clearly identifying *GBA* patient 2 (*GBA*-PSP) (Fig. 2A-B). Using a previously published analysis of the lysosomal proteome (38), we established that 40/72 (56%) of established lysosomal proteins were quantified by examining the sialome with 33/72 (46%) proteins identified by quantifying non-modified peptides (Fig. 2C). Examination of the lysosomal proteome revealed a number of significantly altered lysosomal proteins in *GBA*-PD neurons, including the GWAS-associated lysosomal integral membrane protein 2 (*LIMP2*) (39) and arylsulfatase B (*ARSB*) (40). Additionally, the analysis also confirmed our previous findings of increased levels of glycosylated LAMP1, LAMP2 and cathepsin D in *GBA-*PD iPSC-dopamine neurons (11) (Fig. 2C).

**Fig. 2:**
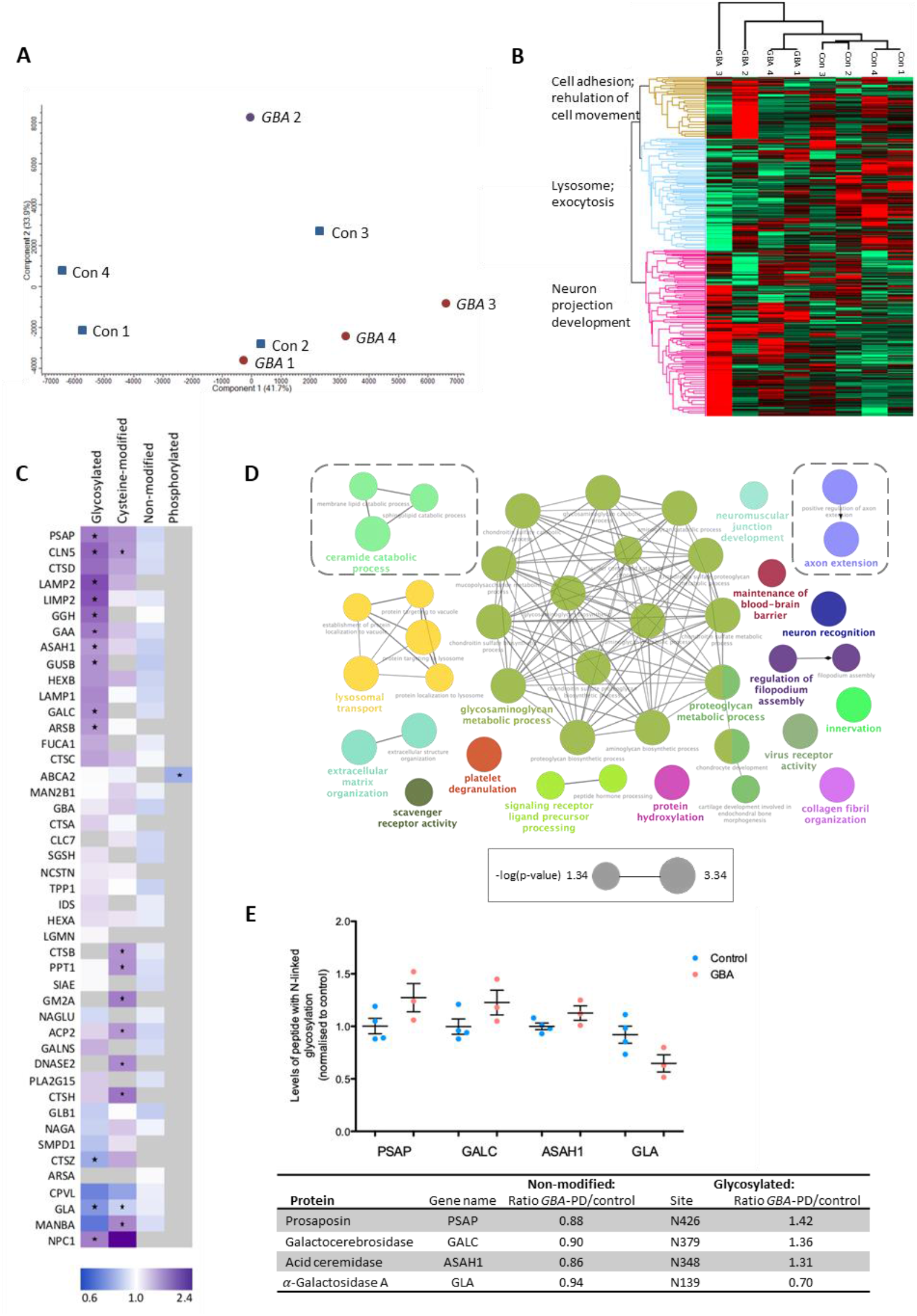
Assessment of glycosylated peptides reveals widespread changes in the lysosomal proteome in *GBA*-PD iPSC-dopamine neurons. A) Principal component analysis (PCA) plot based on glycosylated protein levels in iPSC-dopamine neurons derived from four *GBA*-PD patients (circles; *GBA*-PSP patient in purple) and healthy controls (squares). B) Heatmap of the abundance ratios (*GBA*-PD/control) of all glycosylated proteins identified by the proteomic analysis showing the three main functional clusters that segregate patients and controls. C) Heatmap of the abundance ratios (*GBA*-PD/control) of non-modified protein and PTMs for all established lysosomal proteins identified by the proteomic analysis. Unidentified proteins/PTMs marked with grey. * indicates PTM abundance ratio >1.3 and CV%<30. None of the non-modified proteins were significantly regulated. D) Visualisation of functional GO term networks based on annotations of significantly regulated glycosylated proteins. The node size reflects the enrichment significance of the terms with the leading group term being that of the highest significance. E) Proteomic data from proteins related to ceramide catabolism with graph showing the levels of glycosylated peptides normalised to control and table showing the abundance ratio (*GBA*-PD/control) for the non-modified and glycosylated form including the glycosylation site.

In addition, pathway analysis of the differentially regulated sialated glycoproteome identified a number of significantly-enriched pathways, including ceramide catabolic processes, driven by changes in glycosylation of the PD genome-wide association study (GWAS)-associated genes acid ceramidase (*ASAH1*) (40), galactosylceramidase (*GALC*) (41), prosaposin (*PSAP*) (42) and *α*-Galactosidase A (*GLA*) (43) (Fig. 2D,E). Given the role of GCase in the catabolism of glucosylceramide to glucose and ceramide, the dysregulation of several enzymes involved in ceramide catabolism, which are also GWAS hits for PD, is of particular interest. These data indicate a highly relevant role for the sialated glycoproteome in identifying perturbation in lysosomal biology in neurons. In addition to lysosomal function, differentially regulated sialylated glycoproteins were enriched in processes including neuron projection development and axon extension suggesting a potential novel phenotype in *GBA*-PD (Fig. 2B,D).

### PTMomics identifies tau and mTOR as key mediators in *GBA*-PD pathogenesis

To further understand the biology of *GBA*-PD dopamine neurons, we examined the significantly dysregulated proteins (Fig 3A; Table S2A). Amongst the top upregulated proteins was the microtubule associated protein tau (MAPT), a common GWAS-associated genetic risk locus for PD (44), in addition to a number of other proteins involved in microtubule dynamics, neurite outgrowth and axon extension including tubulin β-2B chain (TUBB2B), MAP1B, MAP6, neuronal cell adhesion molecule (NRCAM), neuromodulin (GAP43), copine-1 (CPNE1) and calponin-2 (CNN2) (Fig. 3A, Fig. S4). Network analysis of the significantly regulated proteins revealed many of these proteins (MAPT, MAP2, MAP6, MAP1B, GAP43) to be part of a larger functional cluster, which also contained a larger number of affected phospho-sites (Fig. 3B). Interestingly, a key protein in this functional cluster was apolipoprotein E (APOE), with levels of APOE demonstrated to be decreased by 0.77-fold in *GBA-*PD (p=0.02) (Fig 3B; Table S2B). Importantly, given the increased prevalence of dementia in *GBA*-PD, dementia-associated *APOE* variants cause decreased APOE levels and increased tau and α-syn pathology (45, 46). Also of relevance to tau function, we identified decreased levels of cysteine-modification of the tau regulator glycogen synthase kinase 3β (GSK3β) at C76, a site which when modified has been shown to inhibit GSK3β activity (47, 48).

**Fig. 3:**
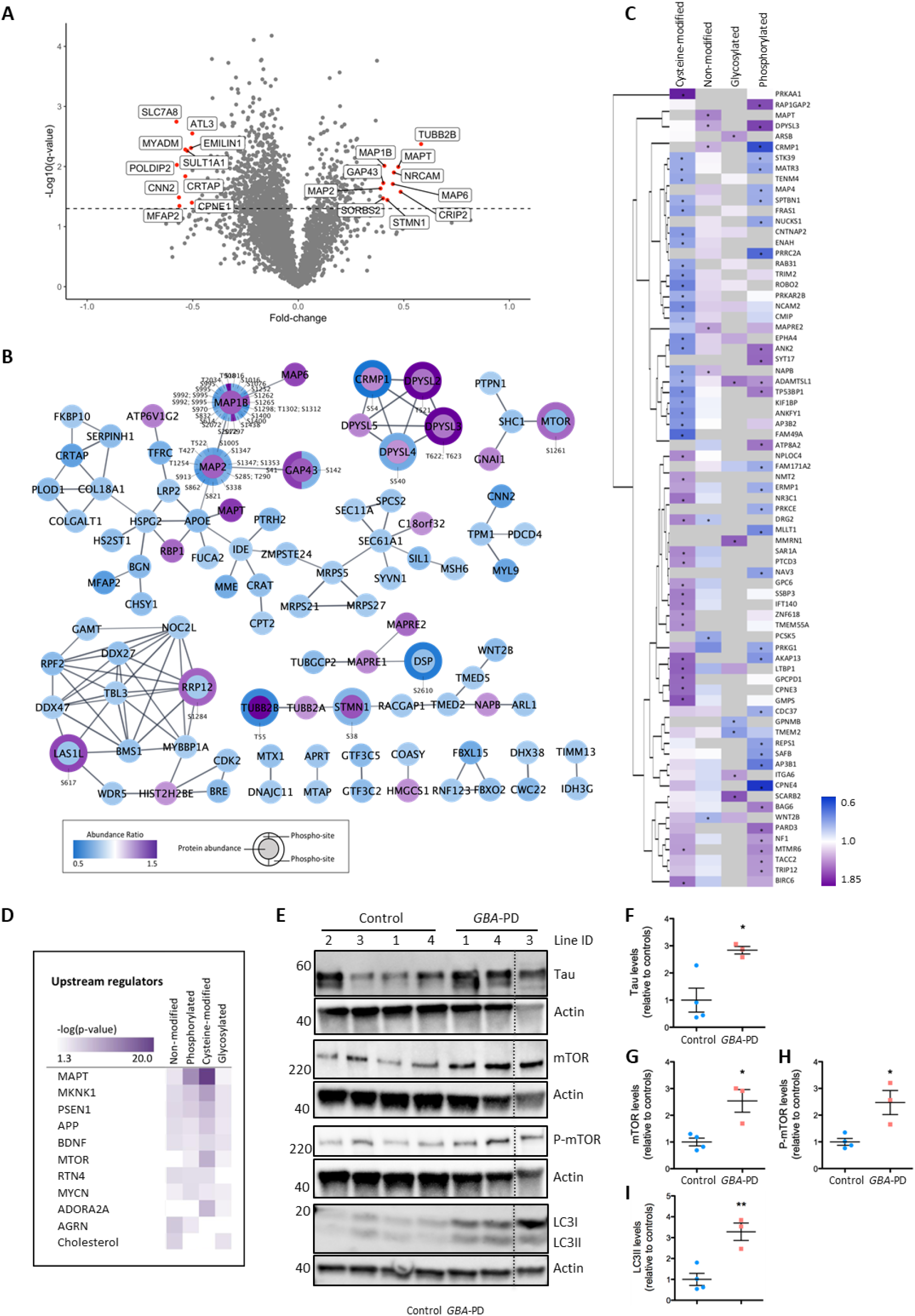
Tau and mTOR are predicted as key regulators of the proteomic changes detected in *GBA-*PD iPSC-dopamine neurons. A) Volcano plot showing the fold change and significance level (-log10 (p-value)) of all non-modified proteins with the dashed line corresponding to a p-value = 0.05. The labelled points represent the ten most highly up/downregulated proteins in GBA-PD patients, relative to control. B) STRING network of connected, significantly regulated non-modified proteins with the circle colour displaying protein abundance ratios (*GBA*-PD/control) and significantly regulated phospho-sites marked by surrounding halo, displaying the phosphorylation abundance ratio (*GBA*-PD/control). C) Heatmap of PD-GWAS proteins with altered protein/PTM levels displaying the abundance ratio (*GBA*-PD/control). Unidentified proteins/PTMs marked with grey. * indicates non-modified protein abundance ratio >1.2 (p-value<0.05) or PTM abundance ratio >1.3 (CV%<30). D) Pathway analysis identifying the predicted upstream regulators most highly enriched across all datasets. Data presented as a heat map of –log(p-value), where increasing colour intensity corresponds to increasing significance in enrichment, as indicated. E-I) Western blotting (E) and quantification of (F) microtubule-associated protein tau (Tau), (G) mammalian target of rapamycin (mTOR), (H) phospho-mTOR and (I) microtubule-associated protein light chain 3 (LC3) levels in *GBA*-PD patient and control neurons included in the mass spectrometry analysis (n = 3-4 iPSC lines). Dashed line in panel E denotes removal of GBA patient 2 from the image. Protein expression levels normalized to β-actin and shown relative to control. Mean ± SEM, * P≤0.05, **P≤0.01 (Student’s *t*-test).

Given the altered levels of PD GWAS targets such as *MAPT* and *ASAH1*, we examined changes in protein levels and PTMs for a list of PD-associated genes compiled from the GWASdb SNP-Disease Associations dataset (49). Of the 756 PD-associated genes, 138 were quantified and 75 (54%) of these were dysregulated in *GBA*-PD neurons (Fig. 3C), corresponding to a significant enrichment of dysregulated proteins encoded by PD-associated genes, compared to the general neuronal proteome (p<0.0001). Particularly, changes in reversible cysteine-modifications were observed on a large number of PD GWAS targets, suggesting differences in the cellular redox environment affecting redox active cysteine redox sites in these proteins (Table S2H).

To understand the biological underpinning of the proteomic changes observed in *GBA*-PD, we performed a directional network analysis to identify the upstream regulators which cause the observed proteomic dysfunction in *GBA*-PD neurons (Fig 3D). Integration of this information across the different datasets highlighted MAPT (tau) as a potential regulator of proteomic dysfunction in *GBA*-PD (Integrated p=0.011), in addition to presenilin 1 (PSEN1), amyloid precursor protein (APP) and mammalian target of rapamycin (mTOR) (Fig. 3D). This strongly suggests that key proteins implicated in PD and AD could be central in mediating *GBA*-related proteomic changes.

Given these data, we confirmed the alterations in the levels of tau in the *GBA*-PD neurons by western blotting (Fig. 3E,F). The proteomic analysis showed a 1.3-fold increase in the active form of the key autophagy regulator, mTOR (phosphorylated at residue S1261) in *GBA-PD* neurons (Fig. 3B, Table S2B-C) (50). This increase in phosphorylated mTOR was confirmed by western blot with a significant increase in levels of mTOR and phospho-mTOR in *GBA*-PD neurons (Fig. 3E,G-H). Given the role of mTOR in regulating multiple pathways, we examined the mTOR downstream targets p70S6K, 4E-BP1 and PKCα. This analysis revealed a significant increase in the ratio of active, phosphorylated 4E-BP1, whereas the other targets were unchanged (Fig. S5A-C). Consistent with previous reports (11) and supporting mTOR activation, we found significantly increased levels of the lipidated form of LC3-II (microtubule-associated protein 1A/1B-light chain 3), an important marker for autophagosome formation, indicating ALP dysfunction (Fig. 3E&I). Together, these data suggest that combined analysis of the proteome and PTMome is able to accurately predict upstream regulators of proteome changes in disease models. Furthermore, we have observed that mTOR activation contributes towards the increased autophagosome accumulation observed in *GBA*-PD neurons.

### PTMomics identifies cytoskeletal organisation and axon extension defects in *GBA*-PD iPSC-dopamine neurons

To further understand the pathways involved in *GBA*-PD neurons, we examined the biological processes which were enriched in the differentially regulated proteins/PTM-peptides. Amongst the non-modified proteins “negative regulation of supramolecular fiber organisation” (p=4.6E-04) and “developmental cell growth” (p=6.1E-04) were some of the most significantly enriched pathways (Fig. 4A, Table S4A), whereas amongst the sialylated glycoproteins “glycosaminoglycan metabolic process” (p=5.2E-09) and “extracellular matrix organisation” (p=6.5E-19) were enriched (Fig. 2D, Table S4B). Amongst the phospho-proteins a high enrichment of RNA-related terms was present including “RNA splicing” (p=8.5E-20) and “regulation of mRNA metabolic process” (p=3.8E-19) (Fig. 4B, Fig. S6, Table S4C). Interestingly, perturbed RNA splicing has recently been suggested as a mediator of PD risk genes (51). In the reversible cysteine-modified group “RNA localisation” (p=6.1E-13) and “ribonucleoprotein complex biogenesis” (p=3.5E-12) were amongst the most highly enriched terms (Fig. 4C, Table S4D).

**Fig. 4:**
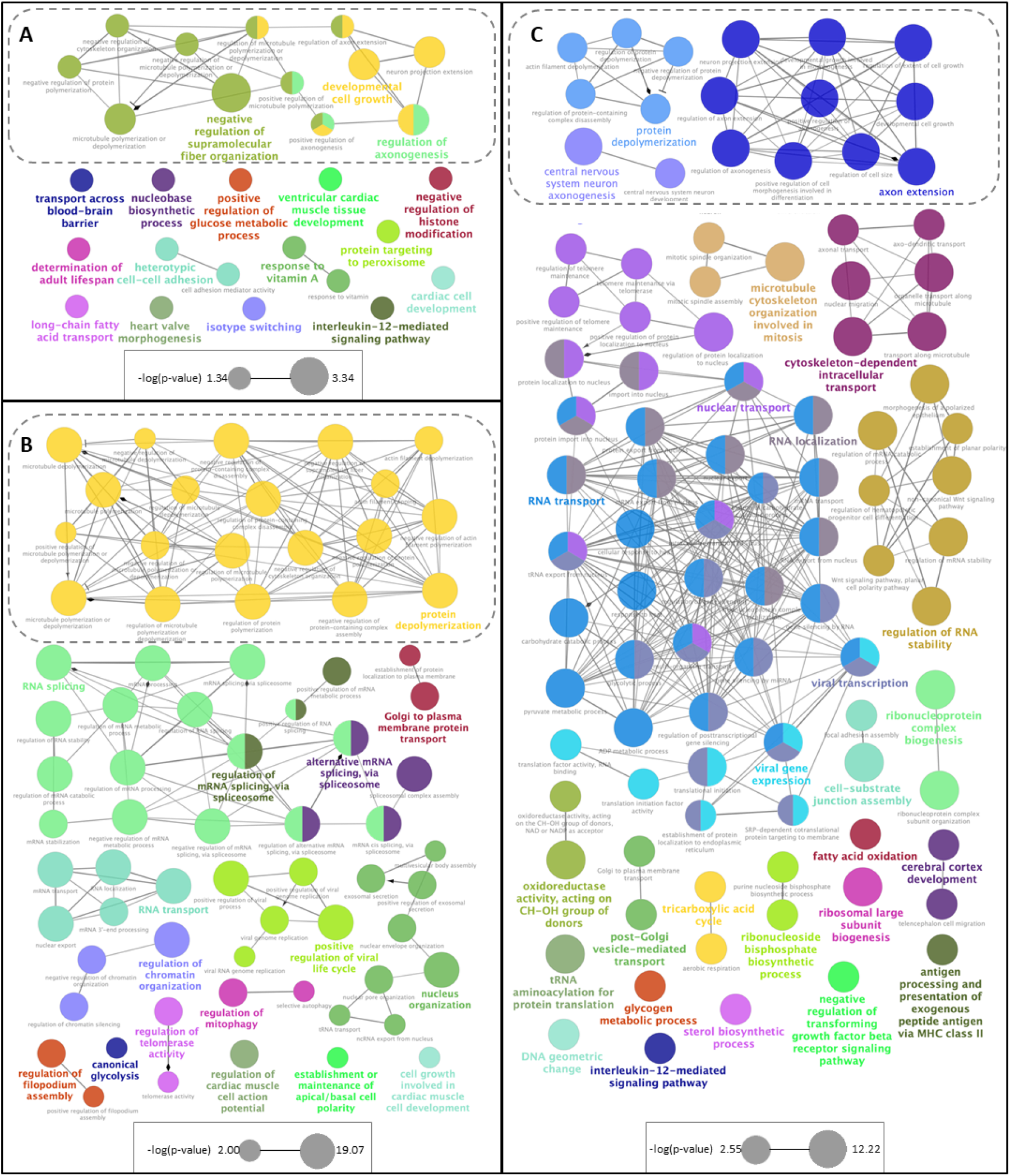
Proteomics and PTMomics indicate cytoskeletal organisation and axon extension defects in *GBA*-PD iPSC-dopamine neurons. A.-C) Visualisation of functional GO term networks based on annotations of significantly regulated (A) non-modified, (B) phosphorylated and (C) cysteine-modified proteins. The node size reflects the enrichment significance of the terms with the leading group term being that of the highest significance. Nodes with similarity in the associated proteins are connected by lines with arrows indicating positive regulation, cropped lines negative regulation and diamonds undefined regulation. Networks related to cytoskeletal organization and axogenesis are marked with dashed squares.

Together, these pathway enrichments suggest that a diverse set of processes are dysregulated in *GBA*-PD neurons. Interestingly, terms related to regulation of cytoskeleton organisation and neuron projection or axogenesis were common to all four groups (Fig. 2D, Fig. 4A-C, Table S4A-D).

### PTMomics reveals tau as a regulator of neurite outgrowth defects in *GBA* patient neurons

Further to this analysis, we performed a pathway analysis on the integrated datasets encompassing significantly dysregulated non-modified and modified proteins. This integrated analysis confirmed “neuritogenesis” (integrated p<0.001) as the most highly enriched pathway when comparing across all four datasets (Fig. 5A).

**Fig. 5:**
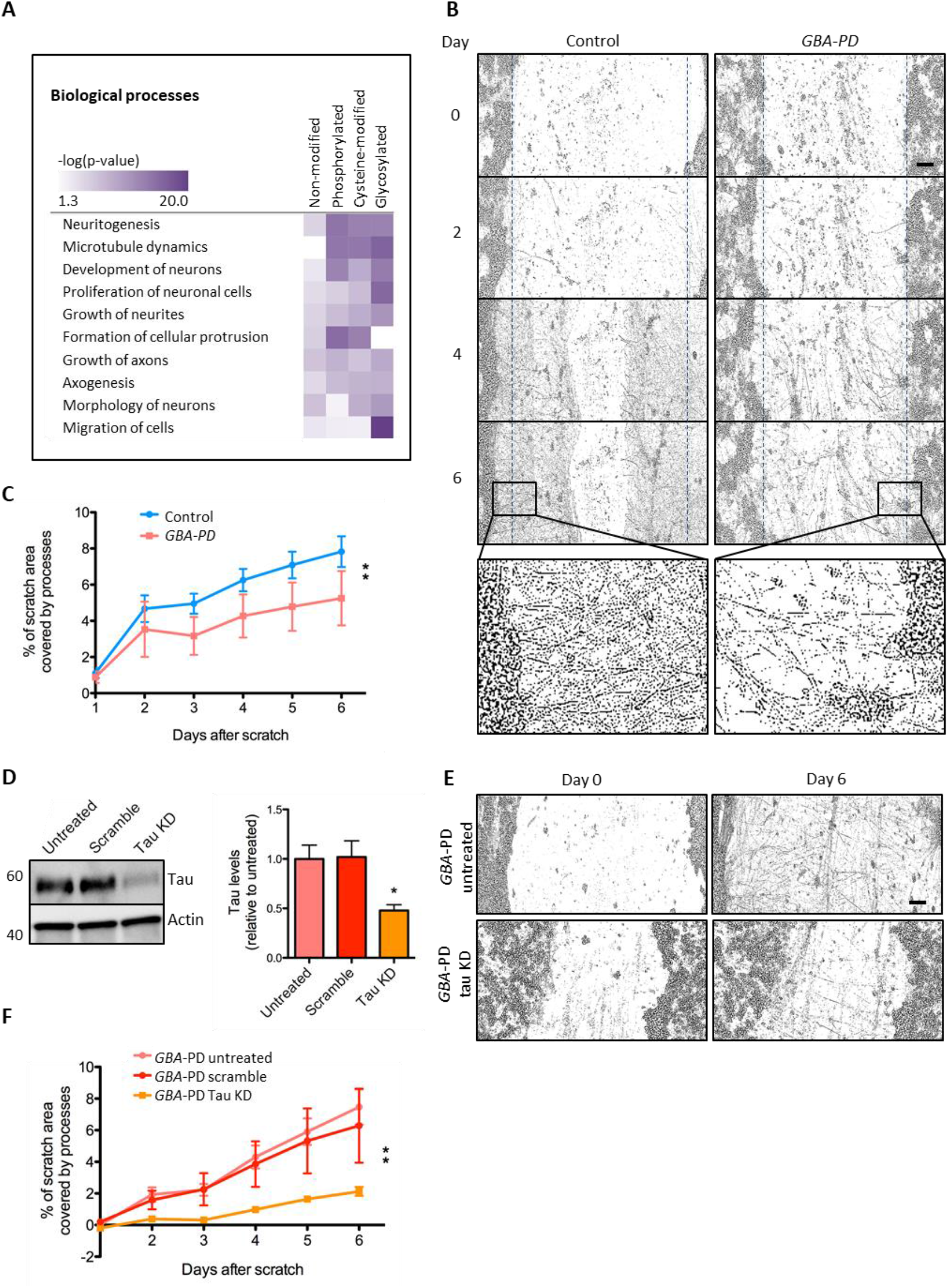
Functional assay confirms defects in neurite outgrowth predicted by pathway analysis. A) Pathway analysis identified the diseases and functions most highly enriched across all datasets. Comparison analysis was performed using Ingenuity Pathway Analysis on the four datasets (non-modified, phosphorylated, glycosylated and cysteine-modified) from the proteomic characterization of *GBA*-PD iPSC-dopamine neurons. Data are presented as a heat map of –log(p-value), where increasing colour intensity corresponds to increasing significance in enrichment. B) Representative bright field images of *GBA*-PD and control iPSC-dopamine neurons from neurite outgrowth assay day 0, 2, 4 and 6 post-scratch. Scratch areas are marked with blue dotted lines. Scale bar = 100 μm. Close-up of neuronal processes in the scratch areas from day 6. Representative close-up of processed binary image used for quantification of neuronal processes in the scratch areas from 6 days post-scratch. C) Quantification of neurite outgrowth as measured by the percentage of the scratch area covered by processes. Values from day 0 subtracted as background. Data from 3 independent differentiations (n = 4-5 iPSC lines per group). Mean ± SEM, **P≤0.01 (Paired Student’s *t*-test). D) Western blotting confirmed lentiviral-mediated shRNA KD of tau in *GBA*-PD iPSC-dopamine neurons (*GBA* 3) and no effect of scrambled shRNA. Expression levels were normalized to β-actin and shown relative to untreated *GBA*-PD iPSC-dopamine neurons (n = 4 technical replicates). Mean ± SEM, *P≤0.05 (Student’s *t*-test). E) Representative bright field images of one line of *GBA*-PD iPSC-dopamine neurons (*GBA* 3) with lentiviral-mediated shRNA tau knock down (KD) and untreated from neurite outgrowth assay day 0, 2, 4 and 6. Scale bar = 100 μm. F) Quantification of neurite outgrowth as measured by the percentage of the scratch area covered by processes over time. Data from *GBA*-PD iPSC-dopamine neurons (*GBA* 3) with shRNA tau KD or a scrambled shRNA control. Values from day 0 subtracted as background (n = 8-12 wells per group). Mean ± SEM, **P≤0.01, (One-way ANOVA, Dunnett’s multiple comparisons).

To investigate the predicted differences in neuritogenesis between *GBA*-PD and control neurons, we utilised an assay for quantification of neurite outgrowth though application of standardized scratches to the monolayer of neurons and measured the growth of neurites into the uncovered scratch area over 6 days (Fig. 5B). Given that very few cells migrated into the scratch area over the course of the assay, the analysis was able to primarily measure neuronal processes extending into the scratch area. Analysis of neurite outgrowth demonstrated a 31% reduction in the ability to regrow into the scratch area in *GBA*-PD neurons (p=0.005), indicating a deficit in neuritogenesis, as predicted by the combined analysis (Fig. 5C).

Given the prominence of tau (MAPT) as an upstream regulator identified in the integrated pathway analysis of *GBA*-PD neurons, we explored the role of the tau in the neurite outgrowth phenotype. We knocked down tau during neuronal maturation using short hairpin RNA (shRNA) and performed the scratch assay. shRNA treatment was confirmed to significantly knockdown total tau levels to around 50% of untreated with the scrambled negative control shRNA having no effect (Fig. 5D; p<0.05). We observed that tau knockdown significantly decreased neurite outgrowth in both a *GBA*-PD patient and a control line, indicating that tau is required for neurite outgrowth in human iPSC-dopamine neurons (Fig. 5E-F, Fig. S7).

### Chaperoning glucocerebrosidase rescues neurite outgrowth defects in *GBA* patient neurons

Given that loss of GCase activity is hypothesised to be a major cause of dysfunction in *GBA-N370S* carriers, we sought to investigate whether rescuing GCase activity could improve the neurite outgrowth phenotype observed in *GBA*-PD neurons. We therefore applied the non-inhibitory small molecule GCase chaperone, NCGC758 (52–54), to day 30 *GBA*-PD iPSC-dopamine neurons and observed a significant, dose-dependent increase in GCase activity four days later (Fig. 6A). Treatment of *GBA*-PD neurons with 15 μM NCGC758 for the duration of the neurite outgrowth assay significantly improved the neurite outgrowth from 43% to 78% of control levels (p=0.016; Fig. 6B), demonstrating the importance of the *GBA* mutation to the observed phenotype and highlighting opportunities for pharmacological rescue.

**Fig. 6:**
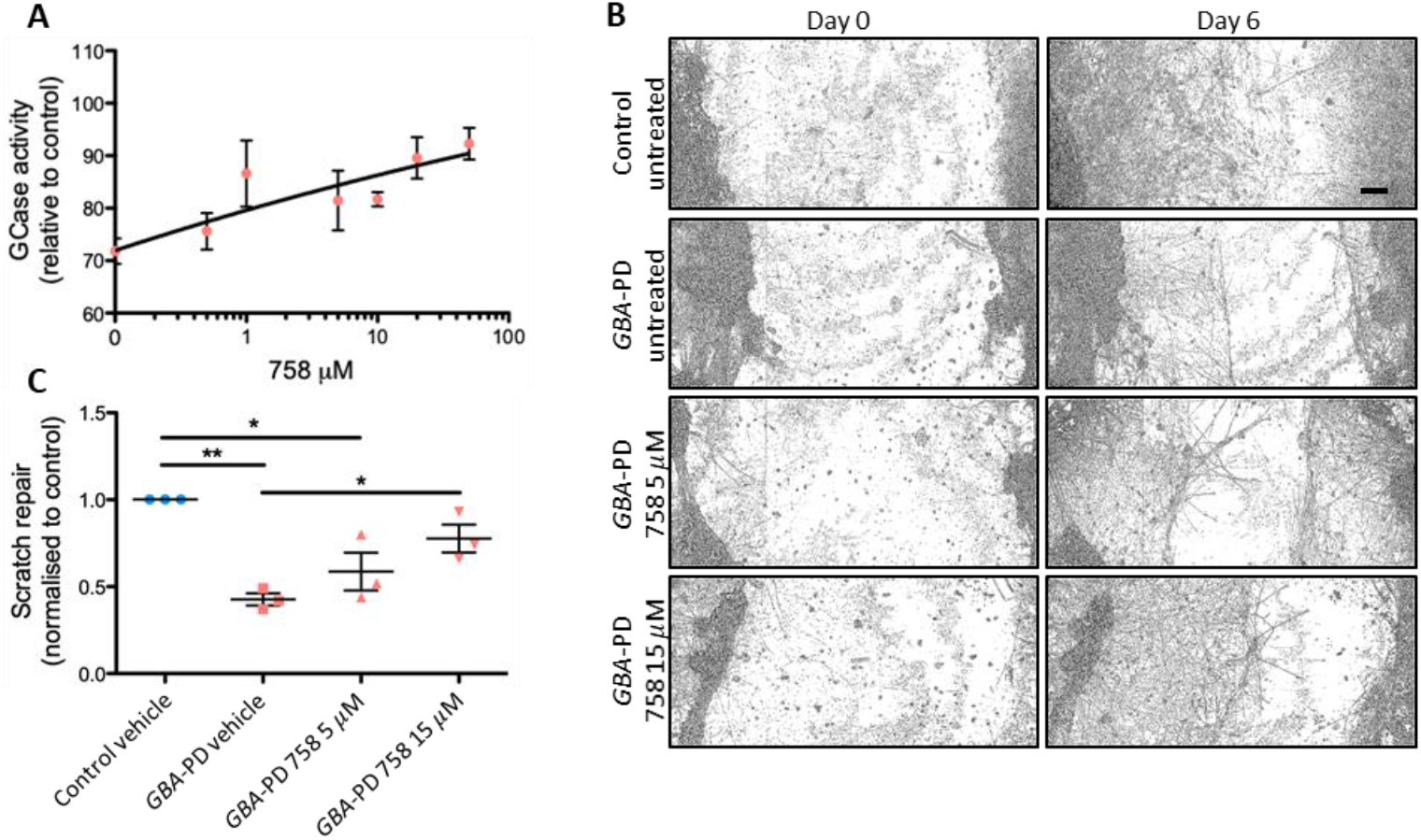
GCase chaperoning rescues neurite outgrowth defects in *GBA*-PD iPSC-dopamine neurons. A) Representative graph of GCase activity in *GBA*-PD iPSC-dopamine neurons treated for 48 h, in response to increasing concentrations of the GCase activator NCGC758 (758), relative to healthy control neurons (n=3-26 wells per condition). Solid line indicates GCase activity in healthy control neurons +/-SEM (dotted line). Datapoints represent mean GCase activity ± SEM (Non-linear regression). B) Representative bright field images of control and *GBA*-PD iPSC-dopamine neurons treated with 5 or 15 μM 758 for 7 days. Scale bar = 100 μm. C. Quantification of scratch repair as measured by the percentage of the scratch area covered by processes on day 6. Values from day 0 subtracted as background. Data from one control and one line of *GBA*-PD iPSC-dopamine neurons (n = 3 independent differentiations). Mean ± SEM, **P≤0.01, (One-way ANOVA, Tukey’s multiple comparisons).

## Discussion

In the present study, we applied a novel post-translational proteomics (TiCPG) workflow to profile the post-translational proteome in human neurons. Mutations in the *GBA* gene are the strongest common genetic risk factor for PD and studies of the effect of *GBA* mutations in human dopaminergic neurons are progressing our knowledge of PD pathogenesis (2, 18, 37). We differentiated *GBA*-*N370S* patient and control iPSC lines using a well-established protocol (55) to generate high numbers of midbrain specific dopaminergic neurons. The simultaneous enrichment and identification of the proteome as well as peptides with phosphorylation, reversible cysteine-modifications and sialylated N-linked glycosylation resulted in an unbiased, global characterisation of PTM levels in human dopaminergic neurons. Moreover, this strategy identified more than 2,000 peptides containing post-translationally modified residues which were dysregulated in *GBA*-PD neurons, making this dataset a resource for research into *GBA*-related pathogenesis. Importantly, this approach utilised small amounts of staring material (100 µg/sample) demonstrating applicability to small-scale cultures of PD patient-derived dopaminergic neurons to identify disease phenotypes and mechanistic targets.

The ability of the workflow to assess sialylated N-linked glycosylation was demonstrated to significantly increase the coverage of lysosomal proteins and gave insight into lysosomal dysfunction in *GBA*-PD, resulting in identification of perturbed levels of several glycosylated lysosomal proteins (11). Sialylated N-linked glycosylation of cathepsin D, LIMP2, LAMP1 and LAMP2 is necessary for their function and transport from the ER via the Golgi to the lysosome and increased lysosomal content is associated with an increase in the glycosylated forms specifically (28, 56). Building on our previous observations of ALP dysfunction in *GBA*-PD lines (11, 20), we identified alterations in the levels of a number of proteins that are encoded by PD GWAS-associated risk genes such as prosaposin (*PSAP*) (42), acid ceramidase (*ASAH1*) (40) and galactocerebrosidase (*GALC*) (41, 57), observations consistent with the known pathways regulated by GCase.

Interestingly, pathway enrichment analysis identified mTOR, an important autophagy regulator, as an upstream target based on PTMomic changes. Although non-modified mTOR was identified as decreased by the mass spectrometry analysis, the phospho-proteomics detected increased mTOR phosphorylation. Western blotting confirmed increased levels of both total and phosphorylated mTOR, suggesting perturbed mTOR signalling which has been implicated in PD pathogenesis based on post-mortem brain studies and cellular models (58). However, the up-regulation of autophagy and lysosomal markers in *GBA* iPSC-dopamine neurons does not suggest mTORC1-mediated autophagy inhibition, and may reflect impaired lysosomal clearance, as previously observed (11). The observation of increased phosphorylation of 4E-BP1, but not p70S6K, suggests increased protein translation via eIF4 (59). This may explain the over-representation of translation as well as RNA transport and splicing in the pathway analysis across the combined datasets. Further studies are needed to determine the consequences of the observed mTOR signalling changes in *GBA-N370S* neurons (58).

The value of large-scale proteomic analysis was also confirmed by the clear separation of PD patients with the *GBA*-*N370S* mutation from controls and the identification of a patient carrying a *GBA-N370S* mutation but with a diagnosis of PSP. The ability of multi-omics analysis of iPSC-dopamine neurons to stratify patients with clinically-overlapping disorders is consistent with our previous work using single-cell RNA-sequencing (RNA-seq) profiling to achieve the same separation (20). Post-hoc comparison of the current study with bulk and single-cell RNA-seq data on *GBA-N370S* patient neurons (20) found a dysregulation of similar genes involved in *MAPT* splicing, microtubule function and formation, neuron projection and axon development.

Although only around 20% of the regulated non-modified proteins had a role in the cytoskeleton, the proteins showing the highest up-regulation were primarily cytoskeletal proteins related to microtubule dynamics and neurite outgrowth. This was also apparent from the pathway analysis which highlighted axon extension across all four datasets. Interestingly, post mortem studies of gene expression changes in early PD have also indicated that the axogenesis pathway is perturbed (60). Our functional analysis confirmed a significant decrease in neurite outgrowth, a phenotype not previously reported in *GBA* iPSC-dopamine neurons. However, iPSC-derived neurons carrying *LRRK2* or *PARK2* mutations or with heterozygous *GBA* knockout also show impaired neurite outgrowth, indicating that this phenotype could be characteristic for PD in general, perhaps even early in the disease development (30, 61, 62).

Tau protein levels were increased in the *GBA*-PD neurons as shown by mass spectrometry and Western blotting. In addition, tau (MAPT) was the top predicted upstream target in the comparison pathway analysis. Tau is not only important in AD pathogenesis, but has been identified by GWAS as a strong risk factor for PD and is found in Lewy bodies co-localising with α-syn (44, 63). In addition, *GBA* mutations are associated with an increased risk of dementia and tau deposition in PD patients (4, 5). To clarify the role of tau in the neurite-outgrowth phenotype, we used shRNA to knock-down tau to around 50% of normal expression levels. This caused a significant inhibition of neurite outgrowth, indicating that tau is required for neurite outgrowth. This confirms the predictions made by the combined analysis and suggests that the role of tau on PD pathology stems from a loss-of-function, perhaps through interaction with α-syn (63).

Interestingly, the neurite outgrowth defect could be rescued using a modulator of GCase activity, NCGC758. This compound is a non-inhibitory GCase chaperone which increases GCase maturation, trafficking and activity, increasing lysosomal GCase activity and reducing α-syn aggregation (37, 52–54). Chaperone rescue by NCGC758 links mutant GCase processing and activity to the neurite outgrowth phenotype in *GBA-*PD patient-derived neurons, adding neuritogenesis to other *GBA-N370S* phenotypes modulated by chaperone treatment (11, 19, 20, 37, 64).

The present proteomic workflow allows for parallel monitoring of three distinct PTMs, in addition to the non-modified peptides observed by traditional proteomics, from a single sample. This represents a substantial improvement on similar workflows (30, 65, 66) and the inclusion of analysis of sialylated glycopeptides resulted in valuable information on alterations in levels of the active/membrane bound forms of numerous lysosomal proteins. The increased coverage of the functional lysosomal proteome in neurons is valuable to disease biology, with significant lysosomal dysfunction found in Parkinson’s, FTD/ALS and Alzheimer’s (67–69). Importantly, this workflow can be performed on as little as 100 μg protein per sample allowing the technique to be performed on small-scale samples, such as brain regions or iPSC-derived neurons.

In conclusion, we have combined *GBA*-PD patient iPSC-dopamine neurons with a comprehensive and unique mass spectrometry methodology allowing the identification of large numbers of dysregulated proteins and PTMs. Integration of the TiCPG datasets led to the discovery of a neurite outgrowth phenotype in iPSC-dopamine neurons from patients carrying the *GBA-N370S* mutation, as well as upstream and downstream modulators of this phenotype, most notably tau. TiCPG analysis of iPSC-derived neurons is thus a powerful method to identify new pathways and phenotypes as well as therapeutic targets and compounds relevant to human neurodegenerative disease.

## Acknowledgements

The work was funded by the Monument Trust Discovery Award (J-1403) from Parkinson’s UK. HB and MM are funded by the Lundbeck Foundation, the Jascha Foundation, the Danish Parkinson Foundation and Innovation Fund Denmark (BrainStem, 4108-00008B). PK is funded by a studentship from the Medical Research Council (MRC) UK. The James Martin Stem Cell Facility, University of Oxford is financially supported by the Wellcome Trust WTISSF121302, and the Oxford Martin School LC0910-004 (SAC), and the MRC Dementias Platform UK Stem Cell Network Capital Equipment and Partnership Awards (SAC and RWM). We thank the High-Throughput Genomics Group at the Wellcome Trust Centre for Human Genetics, Oxford (Funded by Wellcome Trust grant reference 090532/Z/09/Z and MRC Hub grant G0900747 91070) for the generation of Illumina genotyping and transcriptome data. The Villum Center for Bioanalytical Sciences at the University of Southern Denmark is acknowledged for access to state-of-the-art mass spectrometric instrumentation.

## Materials and methods

### Participant recruitment and *GBA-N370S* mutation screening

Participants gave signed informed consent to mutation screening and derivation of iPSC lines from skin biopsies (Ethics committee: National Health Service, Health Research Authority, NRES Committee South Central, Berkshire, UK, REC 10/H0505/71).

PD patients and controls from the Discovery clinical cohort established by the Oxford Parkinson’s Disease Centre were screened for heterozygous *GBA*-*N370S* mutation as earlier described (11). All patients fulfilled the UK Brain Bank diagnostic criteria for clinically probable PD at presentation. Fibroblasts from six *GBA*-*N370S* PD patients and six age and gender-matched healthy controls were reprogrammed to generate iPSC lines (Table S1). Two *GBA* and five control iPSC lines have earlier been characterized (11, 20, 70–72). Detailed characterization of the previously unpublished iPSC lines confirmed their efficient reprogramming, pluripotent state, genome integrity and cell tracking identity (Fig. S8 and S9).

### Culture and reprogramming of primary fibroblasts

Skin punch biopsies were obtained from participants and low passage fibroblast cultures established from these were used for reprogramming either by retroviral delivery or using the CytoTune-iPS Sendai Reprogramming kit (ThermoFisher) as previously described (11, 71). Characterization of previously unpublished iPSC lines was performed on bulk frozen stocks as earlier described (11, 71). Briefly, flow cytometry for pluripotency markers TRA-1-60 (Biolegend #33061, Alexa Fluor 488), and Nanog (Cell Signaling #5448, Alexa Fluor 647) was performed on a FACSCalibur (BD Biosciences). Silencing of retroviral-virus-delivered reprogramming genes was assessed by quantitative real time-PCR (qRT-PCR) using primers published by Takahashi et al. (73), with the following modifications:

pMXsAS3200v2: TTA TCG TCG ACC ACT GTG CTG GCG

mNanog forward primer: GCT CCA TAA CTT CGG GGA GG

For Cytotune 1 and 2 Sendai virus clearance, the following primers were used for standard PCR:

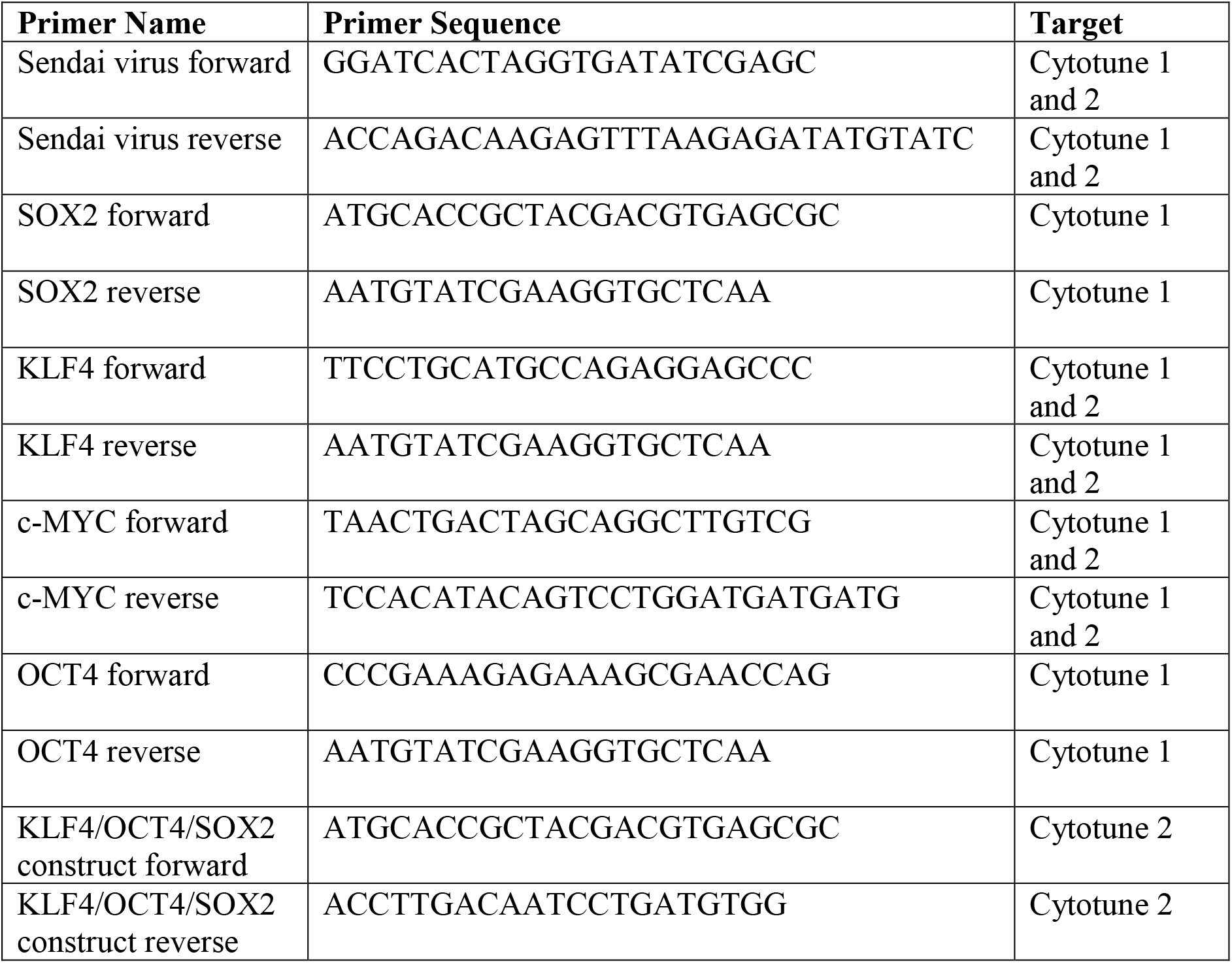

Analysis of pluripotency gene expression profile was performed using the Human-HT-12-v4 expression BeadChip Kit (Illumina) and subsequent Pluritest analysis. Genome integrity and cell identity tracking was assessed using the Human CytoSNP-12v2.1 beadchip array or OmniExpress24 array (Illumina) on genomic DNA generated using the All-Prep kit (Qiagen) and analysis with GenomeStudio and Karyostudio software (Illumina). Illumina SNP datasets and Illumina HT12v4 expression array datasets have been deposited in GEO (awaiting accession number).

### Propagation of iPSCs

Post-thawing iPSCs were propagated as a monolayer on Matrigel-coated (Corning) 6-well plates (Corning) in mTeSR1 medium (Stem Cell Tech.) with 1% penicillin-streptomycin (pen-strep, ThermoFisher), passaged 1:2-3 when confluent using Tryple (ThermoFisher). When thawing or passaging the iPSCs, ROCK inhibitor (Y27632, Bio-Techne) was added to promote survival.

### iPSC differentiation

iPSCs were differentiated using a modified dual-SMAD inhibition protocol as previously described (20, 74). At D20 cells were replated at 3 x 10^5^ cells/cm^2^ and maintained until assays performed at D35.

### Immunofluorescence

Differentiation day 35 cells cultured in 96-well plates (Greiner Bio-One) were fixed for 15 min at room temperature (RT) in 4% (w/v) paraformaldehyde (PFA, Sigma-Aldrich) in distilled phosphate buffered saline (dPBS, ThermoFisher), pH 7.4, and rinsed with dPBS. Cells were permeabilised and unspecific binding blocked with dPBS with calcium and magnesium (dPBS^++^/0.1% Triton (Sigma-Aldrich)/10% donkey serum (Sigma-Aldrich) for 1 hr at RT. Primary antibodies were diluted in dPBS^++^/0.1% (v/v) Triton/1% (v/v) donkey serum and incubated overnight (ON) at 4 °C. The following primary antibody dilutions were applied: rabbit anti-tyrosine hydroxylase (TH, Millipore #152) 1:500, goat anti-FOXA2 (R&D #AF2400) 1:250, mouse anti-β-III-Tubulin (TUJ1, Biolegend #801202) 1:500.

Cultures were rinsed in PBS and incubated with donkey anti-rabbit AF488 (ThermoFisher #A21206) 1:1000, donkey anti-goat AF647 (ThermoFisher #A21447) 1:500 and donkey anti-mouse AF555 (Invitrogen #A31570) 1:1000 in dPBS^++^/0.1% Triton/1% donkey serum for 1 hr at RT. Cell nuclei were counterstained with 10 μM 4“,6-diamidino-2-phenylindole dihydrochloride (DAPI, Sigma-Aldrich) in dPBS^++^. Fluorescence images were acquired on the Opera Phenix High-Content Screening Confocal microscope (Perkin-Elmer).

### Glucocerebrocidase (GCase) activity assay

GCase activity was measured by cleavage of 4-methylumbelliferyl-β-D-glucopyranoside (4-MUG) to 4-methylumbelliferone, as previously described (31). Briefly, cell pellets were sonicated at 10 amp for 10 sec in citrate phosphate buffer pH 5.4 consisting of 0.1 M citric acid (Sigma-Aldrich) and 0.2 M dibasic sodium phosphate (Sigma-Aldrich) with 0.25% (v/v) Triton X and 0.25 % (w/v) taurocholic acid (Sigma-Aldrich). Samples were centrifuged at 800 g for 5 min at 4 °C and supernatant collected. Following protein determination equal amounts of protein from each sample were diluted in citrate phosphate buffer in quadruplicates. One replicate of each sample was treated with 1 mM conduritol B epoxide (CBE, Calbiochem) for 10 min before all samples were incubated for 1 hr with 2.5 mM of the fluorescent GCase substrate methylumbillifery β-D-glucopyranoside (4MUG, Sigma-Aldrich) at 37 °C in the dark. The reaction was quenched with 1 M glycine buffer (Sigma-Aldrich) pH 10.8 and the fluorescent levels analysed on the PHERAStar FSX plate reader (BMG Labtech). Values from CBE-treated wells were subtracted as background.

### Western blotting procedure

Cell pellets were lysed in RIPA buffer (50 mM Tris-hydrochloride (Sigma-Aldrich), 150 mM sodium chloride (Sigma-Aldrich), 1% Tergitol-type NP-40 (Sigma-Aldrich), 0.5% sodium cholate hydrate (Sigma-Aldrich), and 0.1% SDS (Sigma-Aldrich), pH 8) containing phosphatase (PhosSTOP tablets, Roche) and protease inhibitor (Complete Tablets, Roche) and sonicated for 10 secs at 10 amplitude microns on ice. Protein concentrations were determined with bicinchoninic acid assay (BCA, Pierce) and equal amounts of protein were separated by SDS-PAGE. Membranes were probed overnight with primary antibodies as follows: mouse anti-β-Actin-HRP (Abcam #49900) 1:50000, rabbit anti-LC3 (Sigma-Aldrich #L7543) 1:1000, mouse anti-Tau (Neomarkers #ms-247-P) 1:1000, mouse anti-mTOR (Cell Sig. #45175) 1:1000, rabbit anti-phospho-mTOR (Ser2448) (Cell Sig. #2971) 1:100, rabbit p70S6K (Cell Sig. #2708) 1:1000, mouse anti-phospho-p70S6K (Thr389) (Cell Sig. #9206) 1:100, rabbit anti-4E-BP1 (Cell Sig. #4923) 1:1000, rabbit anti-phospho-4E-BP1 (Thr37/46) (Cell Sig. #2855) 1:100, rabbit anti-PKCα (Cell Sig. #2056) 1:1000, and rabbit anti-phospho-PKCa (Ser657) (ThermoFisher #PA5-78124) 1:500. Representative blots of all antibodies shown in Fig. S10.

Membranes were subsequently probed with horseradish peroxidase (HRP)-conjugated goat anti-mouse or -rabbit IgG (Bio-Rad), diluted 1:5000 in 5% (w/v) skim milk in TBST, for one hr and visualised using ECL (Millipore #P90720) on a ChemiDoc Touch Imaging system (Bio-Rad). The optical density of each band was quantified using Image Lab software (Bio-Rad).

### Mass spectrometry sample collection and protein isolation

Differentiation day 35 neurons from four *GBA* and four control iPSC lines were collected on ice in phosphate-buffered saline (PBS, ThermoFisher) with protease-(Complete tablets, Roche) and phosphatase inhibitors (PhosSTOP tablets, Roche). Samples were sonicated for two times ten secs at 50% amplitude on ice and incubated for 30 min at RT in lysis buffer consisting of 6 M urea (Sigma-Aldrich), 2 M thiourea (Sigma-Aldrich), 20 mg/ml sodium dodecyl sulfate (SDS, GE Healthcare), 40 nM N-ethylmaleimide (NEM, for alkylation of free cysteines, Sigma-Aldrich) and protease inhibitor.

### Reduction and enzymatic digestion

Following methanol-chloroform precipitation (Sigma-Aldrich), the proteins were denatured and reduced in 6 M urea, 2 M thiourea and 10 mM TCEP (ThermoFisher) at RT. After vortexing, the samples were incubated at RT for 2 hrs with 1 μl endoproteinase Lys-C (Wako). The samples were diluted 10 times in 20 mM TCEP in 20 mM triethylammonium bicarbonate (TEAB, Sigma-Aldrich) buffer, pH 7.5 and sonicated for two times ten seconds at 50% amplitude on ice. Digestion with 1 μg trypsin (Sigma-Aldrich) per 50 μg peptide was done ON at RT.

### Desalting and iTRAQ labelling

The samples were acidified with 0.1% trifluoroacidic acid (TFA, Sigma-Aldrich) and desalted using two self-made P200-tip-based columns per sample. A small plug of C_18_ material from a 3M Empore^TM^ disk (Sigma-Aldrich) was inserted in the constricted end of a P200 pipette tip and 1.5 cm of the tip was packed with reversed-phase resin material consisting of a 1:1 mix of Poros 50 R2 (Applied Biosystems) and Oligo R3 (Applied Biosystems) dissolved in 100% acetonitrile (ACN, Sigma-Aldrich). The acidified samples were loaded onto the first micro-columns and washed with 0.1% TFA. The peptides were eluted using 60% ACN/0.1% TFA. This process was repeated with the second column before the samples were dried by speed vacuum centrifugation.

100 μg of each sample were labelled with Isobaric Tags for Relative and Absolute Quantification (iTRAQ) Eight-plex Isobaric label Reagents (AB Sciex) according to the manufacturer’s instructions. Efficient labeling was confirmed by MALDI, the ratios adjusted, the labeled peptides mixed 1:1:1:1:1:1 and dried.

### Enrichment of phosphorylated, sialylated glyco- and cysteine-modified peptides

Phospho-peptide enrichment and fractionation were essentially performed as earlier described (75). Peptides were dissolved in 80% ACN/5%TFA with 1 M glycolic acid (Sigma-Aldrich) and incubated with 0.6 mg titanium dioxide (TiO_2_) beads (Titansphere 10 µm, GL Sciences) per 100 mg peptide for 30 min at RT with vigorous shaking. The beads were centrifuged briefly and the supernatant transferred to a new tube with 0.3 mg TiO_2_ beads per 100 mg peptide. After 15 min incubation at RT with vigorous shaking and a brief centrifugation the supernatant was collected. The beads were subsequently washed with 80% ACN/1% TFA and 10% ACN/0.1% TFA. The supernatant with the unbound TiO_2_ fraction and the washing fractions, both containing the non-phosphorylated peptides, were combined and dried for further processing (see below). The phosphorylated peptides were eluted from the beads by incubation with 1.5% ammonium hydroxide solution (Sigma-Aldrich), pH 11.3, at RT with vigorous shaking. The beads were spun down and the supernatant passed through C_8_ material from a 3M Empore^TM^ disk (Sigma-Aldrich). Any remaining peptides were eluted from the filter with 30% ACN and all peptide samples were dried.

The dried phosphorylated peptides were dissolved in 20 mM TEAB, reduced with 1 M dithiothreitol (DTT, Sigma-Aldrich) for 30 min at RT and alkylated with 1 M NEM for 30 min at RT in the dark. To separate out sialylated glyco-peptides that also bind to the TiO_2_ beads, the sample was deglycosylated with N-glycosidase F (Biolabs) and Sialidase A (Prozyme) at 37°C ON. The sample was dried, resuspended in SIMAC loading buffer (50% ACN/0.1% TFA) and incubated with PhosSelect IMAC beads (Sigma #P9740), pre-equilibrated in SIMAC loading buffer, for 30 min at RT with gentle shaking. The beads were washed in SIMAC loading buffer before mono-phosphorylated and deglycosylated peptides were eluted with 20% ACN/1% TFA, combined with the washing fraction and dried. The multi-phosphorylated peptides were then eluted with 1.5% ammonium hydroxide solution, pH 11.3, and dried. The mono-phosphorylated and deglycosylated peptides were separated by adjusting the sample to 70% ACN/2% TFA and repeating the TiO_2_ bead enrichment as described above (washing with 50% ACN/0.1%TFA). Deglycosylated peptides were collected in the supernatant.

The dried non-phosphorylated sample was desalted on R2/R3 columns as described above and dissolved in 50 mM TEAB and 20 mM TCEP, pH 7, for 1 hr. The sample was incubated with 10 mM cysteine-specific phosphonate adaptable tag (CysPAT), synthesized as previously described (76), for 1 hr at RT in the dark with gentle shaking. The CysPAT reacts with reversibly modified cysteines and the phosphonate group allows for the subsequent isolation of CysPAT-labeled peptides from non-modified peptides using TiO_2_ bead enrichment as earlier detailed.

All the eluates were dried and desalted on homemade columns as described previously, using only R3 material for the mono- and multi-phosphorylated peptides and R2/R3 material for deglycosylated, cysteine- and non-modified peptides.

### Hydrophobic interaction liquid chromatography (HILIC) and high pH fractionation

Mono-phosphorylated-, deglycosylated-, CysPAT-labeled- and non-modified peptides were fractionated to reduce sample complexity using HILIC as described previously (75). The multi-phosphorylated sample was not fractionated, but run directly by LC-ESI-MS/MS. The non-modified sample, expected to contain the highest concentration of peptides, was first diluted in 0.1% TFA and approximately 50 mg peptide was fractioned. All mono-phosphorylated-, deglycosylated- and CysPAT samples were fractionated. The samples were dissolved in 90% ACN, 0.1% TFA (solvent B) by adding 10% TFA followed by water and finally ACN slowly in order to prevent peptide precipitation. Samples were loaded onto an in-house packed TSKgel Amide-80 (Tosoh Bioscience) micro-capillary column (450 mm OD x 320 mm ID x 17 cm) using an Agilent 1200 Series HPLC (Agilent). Peptides were separated using a gradient from 100–60% solvent B (A = 0.1% TFA) running for 30 min at a flow-rate of 6 ml/min. Fractions were collected every 1 min and combined into 8-12 final fractions based on the UV chromatogram and subsequently dried by vacuum centrifugation.

To increase the coverage, high pH fractionation was also performed using approximately 50 mg peptide of the non-modified peptide sample. Briefly, the sample was dissolved in 1% ammonium hydroxide (NH_3_, Sigma-Aldrich), pH 11, and loaded on a R2/R3 column equilibrated with 0.1% NH_3_. The peptides were eluted in a stepwise fashion using a gradient of 5%-60% ACN/0.1% NH_3_. All fractions were dried by vacuum centrifugation and stored at −20 °C.

### Reversed-phase nanoLC-ESI-MS/MS

The samples were resuspended in 0.1% formic acid (FA) and loaded onto an EASY-nLC system (Thermo Scientific). The samples were loaded onto a two-column system containing a 3 cm pre-column and a 17 cm column both consisting of fused silica capillary (75 μm inner diameter) packed with ReproSil–Pur C18 AQ 3 μm reversed-phase material (Dr. Maisch). The peptides were eluted with an organic solvent gradient from 100% phase A (0.1% FA) to 34% phase B (95% ACN, 0.1% FA) at a constant flowrate of 250 nL/min. Depending on the samples based on the HILIC, the gradient was from 1 to 30% solvent B in 60 min or 90 min, 30% to 50% solvent B in 10min, 50%-100% solvent B in 5 min and 8 min at 100% solvent B.

The nLC was online connected to a QExactive HF Mass Spectrometer (Thermo Scientific) operated at positive ion mode with data-dependent acquisition. The Orbitrap acquired the full MS scan with an automatic gain control (AGC) target value of 3×10^6^ ions and a maximum fill time of 100ms. Each MS scan was acquired at high-resolution (120,000 full width half maximum (FWHM)) at m/z 200 in the Orbitrap with a mass range of 400-1400 Da. The 12 most abundant peptide ions were selected from the MS for higher energy collision-induced dissociation (HCD) fragmentation (collision energy: 34V). Fragmentation was performed at high resolution (60,000 FWHM) for a target of 1×10^5^ and a maximum injection time of 60ms using an isolation window of 1.2 m/z and a dynamic exclusion. All raw data were viewed in Thermo Xcalibur v3.0.

### Mass spectrometry data analysis

The raw data were processes using Proteome Discoverer (v2.1, ThermoFisher) and searched against the Swissprot human database using an in-house Mascot server (v2.3, Matrix Science Ltd.) and the Sequest HT search engine.

The mass spectrometry proteomics data have been deposited to the ProteomeXchange Consortium via the PRIDE(77) partner repository with the dataset identifier PXD026691.

Database searches were performed with the following parameters: precursor mass tolerance of 10 ppm, fragment mass tolerance of 0.02 Da (HCD fragmentation), TMT 6-plex (Lys and N-terminal) as fixed modifications and a maximum of 2 missed cleavages for trypsin. Variable modifications were NEM on Cys and N-terminal acetylation along with phosphorylation of Ser/Thr/Tyr, deamidation of Asn and N-Succinimidyl iodoacetate (SIA) on Cys for the phosphorylated, deglycosylated and CysPAT-modified groups, respectively. Only peptides with up to a q-value of 0.01 (Percolator), Mascot rank 1 and cut-off value of Mascot score > 15 were considered for further analysis. Only proteins with more than one unique peptide were considered for further analysis in the non-modified group. Subsequently, proteins which were at least 1.2-fold up-or down-regulated in the *GBA* patient neurons were selected and Student’s t-test with Benjamini-Hochbergs correction for multiple testing (FDR 0.1) was applied with a p-value cut-off of 0.05.

Peptides with PTMs which 1. could be normalised to the level of the corresponding non-modified protein, 2. were at least 1.3-fold up-or down-regulated in the *GBA* patient neurons and 3. had a coefficient of variation (CV) of 30% or less were selected for further analysis. Furthermore, peptides with N-linked glycosylation (NxS/T/C motif) were manually sorted based on information from UniProt (78) on known glycosylation and cellular localisation (Golgi/endosome/lysosome/membrane/extracellular) to exclude spontaneous deamidations.

The pathway analysis was performed using the Ingenuity Pathway Analysis software (IPA, QIAGEN) on the four datasets; the differentially regulated non-modified-, phosphorylated-, glycosylated- and cysteine-modified proteins. Direct and indirect relationships, either experimentally observed or highly predicted based on data from CNS tissue or cell lines, were included. The IPA comparison analysis was used to combine results from all four datasets and rank the most significantly enriched pathways according to their IPA calculated p-values. An integrated p-value was calculating based on the mean of the p-values from each dataset.

The Cytoscape StringApp (3.8.2) was used to visualize networks of differentially expressed proteins and PTMs using the Omics Visualizer app with a confidence score cut-off of 0.7 (79). Furthermore, enrichment analysis was performed focusing on Reactome pathways in the STRING Enrichment app. Hierarchical clustering of the 500 most abundant proteins was computed using the ClusterMaker app analysing the pairwise average linkage with Euclidean distance as distance metric and results shown as a heatmap. The ClueGO (2.5.6) Cytoscape plug-in was applied to visualize, which functionally grouped Gene Ontology (GO) terms in the category Biological process the differentially expressed proteins in the four datasets were annotated to (80). The Homo sapiens (9606) marker set was applied with the following settings: GO tree levels of minimum 3 and maximum 8 and a Kappa score threshold of 0.4. For GO term selection a minimum of 3 genes and 4% of genes were used for the non-modified, whereas a minimum of 4/5/10 genes and 5/8/15% were used for the larger lists of glycosylated proteins, phosphorylated and cysteine-modified proteins, respectively.

PCA plots and heatmaps for PTM datasets were generated using Perseus (81). Heatmaps were generated from z-score median normalised data and were clustered using Euclidean distance and k-means clustering. Subsequent categorical enrichment was performed using g:Profiler (82) and significant GO molecular function/cellular component/biological process and KEGG terms aggregated to provide an overview of cluster function/compartment.

Enrichment of PD-associated genes was calculated using Fisher’s exact test and based on the number of proteins identified in one or more of the datasets (Table S2A,C,E,G) and the number of regulated proteins showing protein and/or PTM level changes (Table S2B,D,F,H) compared to the number of identified and regulated PD-associated proteins.

### Neurite outgrowth assay

Scratches were applied to differentiation day 30 neurons across the centre of each well in a 96-well plate using a 10 μl pipette tip followed by replacement of the neuronal maturation medium. Cells were treated with NCGC00188758 (NCGC758, Sigma-Aldrich #5316600001) after the scratch was applied and media +/- NCGC758 was replaced every 48h. The scratch area was imaged daily for a week using the Opera Phenix High-Content Screening Confocal microscope (Perkin-Elmer). Images were analysed using ImageJ by sharpening and converting images to binary, marking the scratch area and for each day calculating the area of the scratch covered by processes (particle size: 3-1000, circularity: 0.0-0.8) (83). Values from day 30 were subtracted as background.

Lenti-virus construction and production of shRNA targeting MAPT exons 12-13 (constitutive exons for targeting total MAPT) and no known RefSeq transcript (scrambled, non-targeting shRNA) was performed as previously described (84). IPSC-derived neuronal precursors were transduced with lentiviral particles encoding shRNAs targeting MAPT or scrambled on differentiation day 20 following replating.

### Statistical analysis

Analysis was performed in Graphpad Prism version 5.0 (GraphPad Software, USA) using two-tailed paired or unpaired Student’s *t*-tests, one sample *t*-test, Fisher’s exact test and one-way ANOVA with Dunnett’s multiple comparison, where appropriate. Results are expressed as mean ± SEM, and p-values ≤ 0,05 were considered statistically significant.

## Supplementary Information

**Fig. S1:**
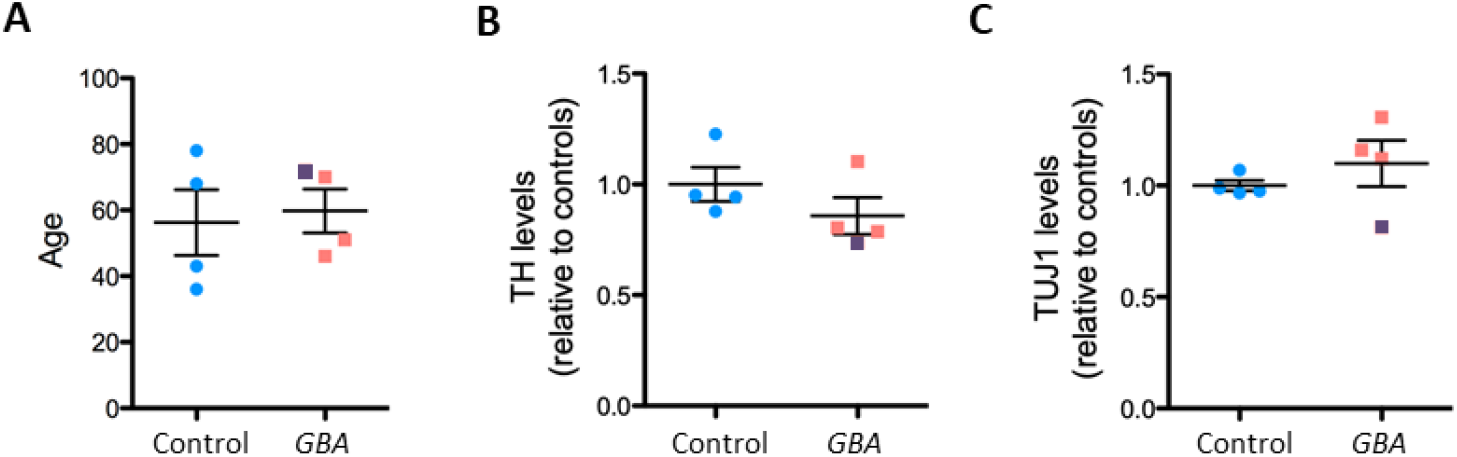
Characterisation of iPSC lines and iPSC-dopamine neurons applied in the mass spectrometry analysis. A) Age of the control individuals and *GBA* patients included in the proteomic analysis (n = 4 iPSC lines per group). One *GBA* patient (*GBA* 2, shown in purple) was later discovered to be L-DOPA unresponsive and diagnosed with PSP. B-C) Mass spectrometry quantification of (B) TH levels (identified with 17 unique peptides, 34.6% coverage) and (C) TUJ1 levels (identified with 14 unique peptides, 75.6% coverage) in *GBA* patient and control neurons included in mass spectrometry analysis shown relative to control neurons (n = 4 iPSC lines per group). *GBA* 2 (*GBA*-PSP) in purple. Mean ± SEM (Student’s *t*-test).

**Fig. S2:**
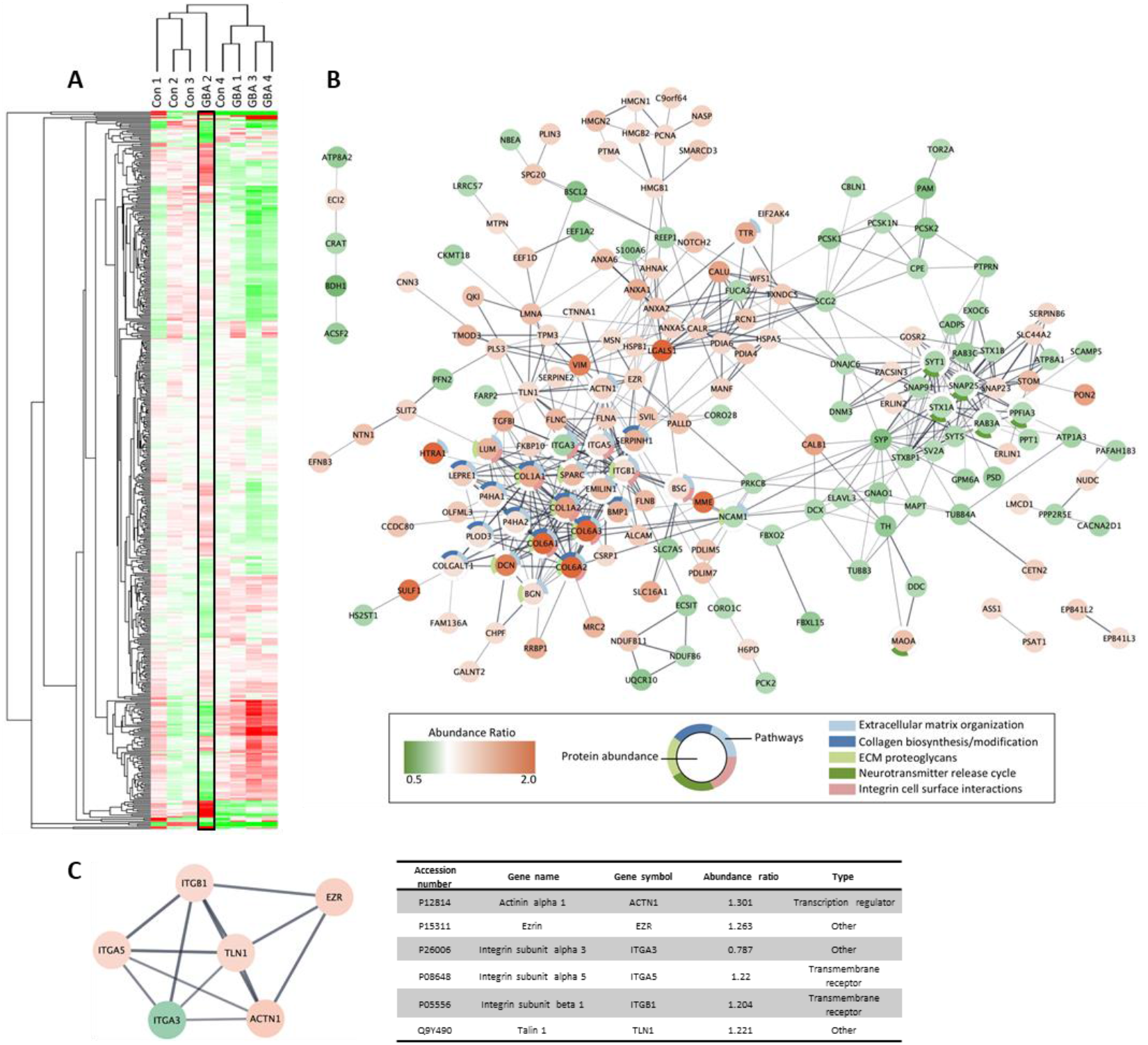
Pathway analysis comparing *GBA* 2 vs. control. A) Heatmap of the abundance ratios (*GBA*/control) of the 500 most abundant proteins identified by the mass spectrometry analysis with data for *GBA* 2 (iPSC line 834) highlighted. B) STRING network of significantly regulated non-modified proteins with the circle colour displaying non-modified protein abundance ratios (*GBA* 2/control) and surrounding halo marking proteins belonging to one or more of the five Reactome pathways with the lowest FDR value based on STRING enrichment analysis. C) Pathway analysis of the significantly regulated non-modified proteins (*GBA* 2/control) identified “Calpain protease regulation of cellular mechanics” as the top canonical pathway based on enticement of the proteins shown in the network/table.

**Fig. S3:**
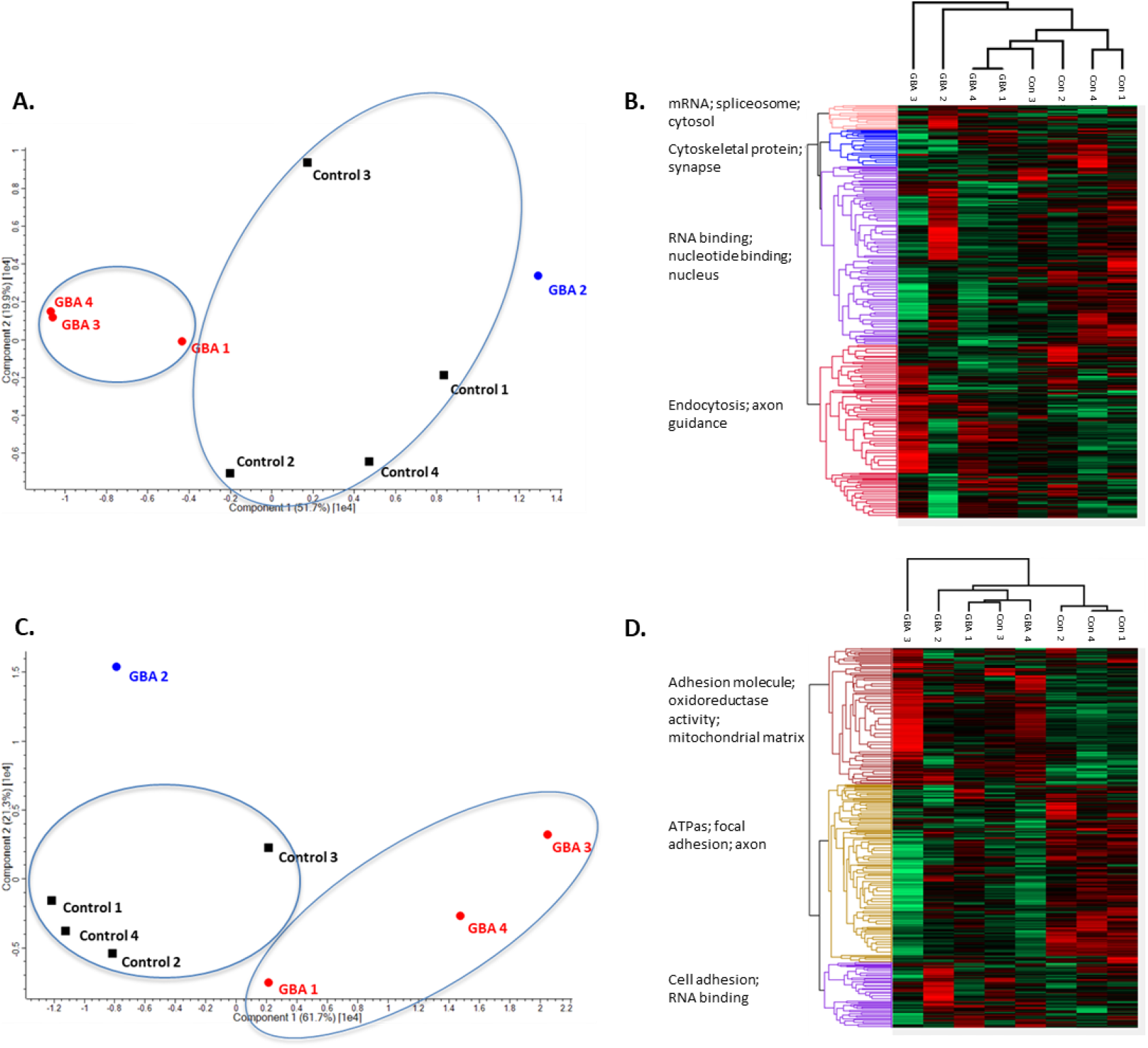
Segregation of *GBA*-PD, *GBA*-PSP and control lines based on PTMs. A.-D) PCA plot based on (A) phosphorylated and (C) cysteine-modified peptide levels in G*BA* patient (circles, *GBA* 2 (PSP) in purple) and healthy control neurons (squares). Heatmaps of the abundance ratios (*GBA*/control) of (B) phosphorylated and (D) cysteine-modified peptides identified by proteomic analysis showing the main functional/subcellular compartment clusters in the dataset.

**Fig. S4:**
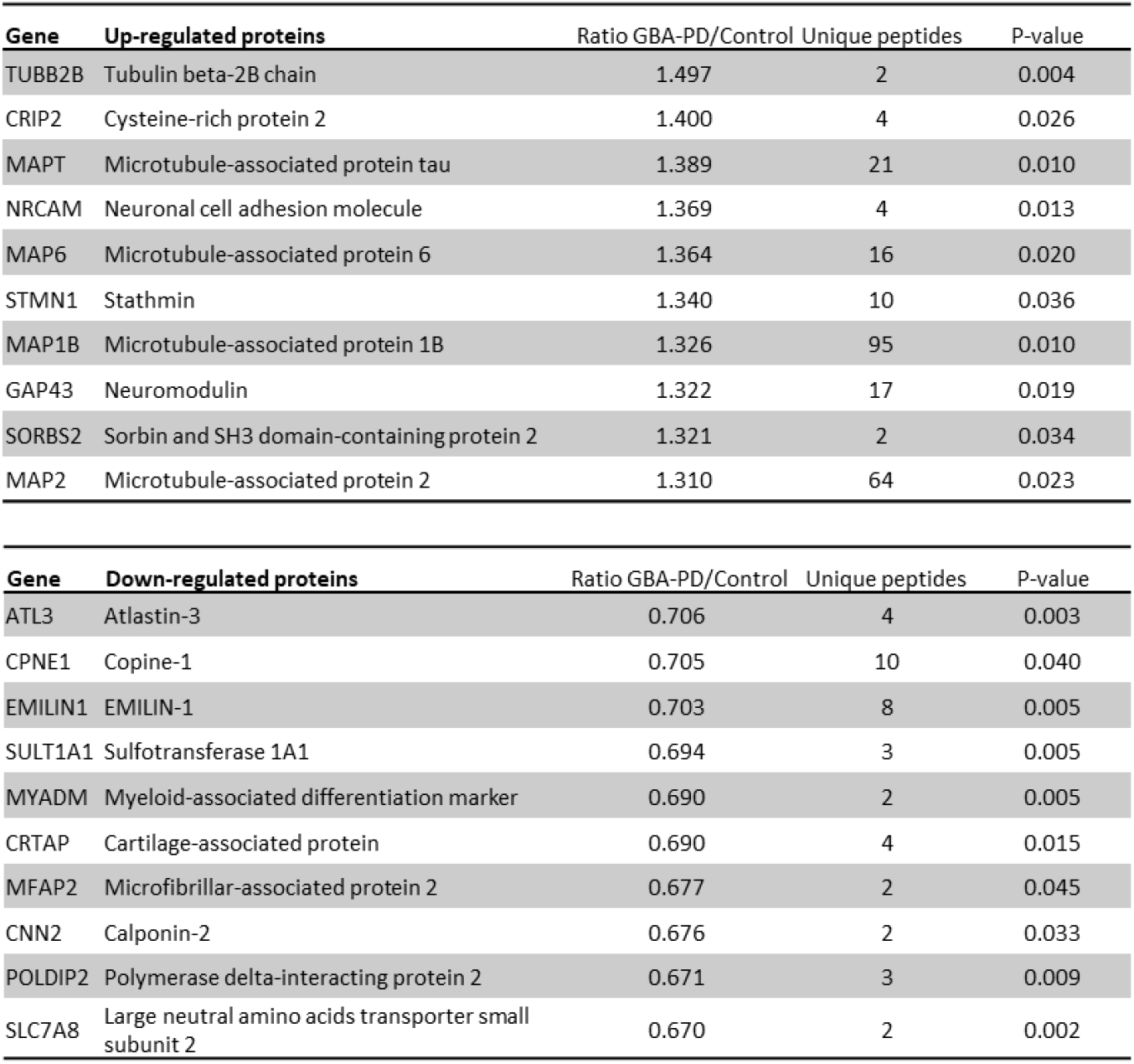
The most up- and down-regulated proteins in *GBA* patient neurons. The non-modified proteins identified by the mass spectrometry analysis as most up- and down-regulated in the *GBA*-PD patient neurons. Expression levels shown as ratio of *GBA*-PD relative to control with p-value and number of unique peptides used for protein identification.

**Fig. S5:**
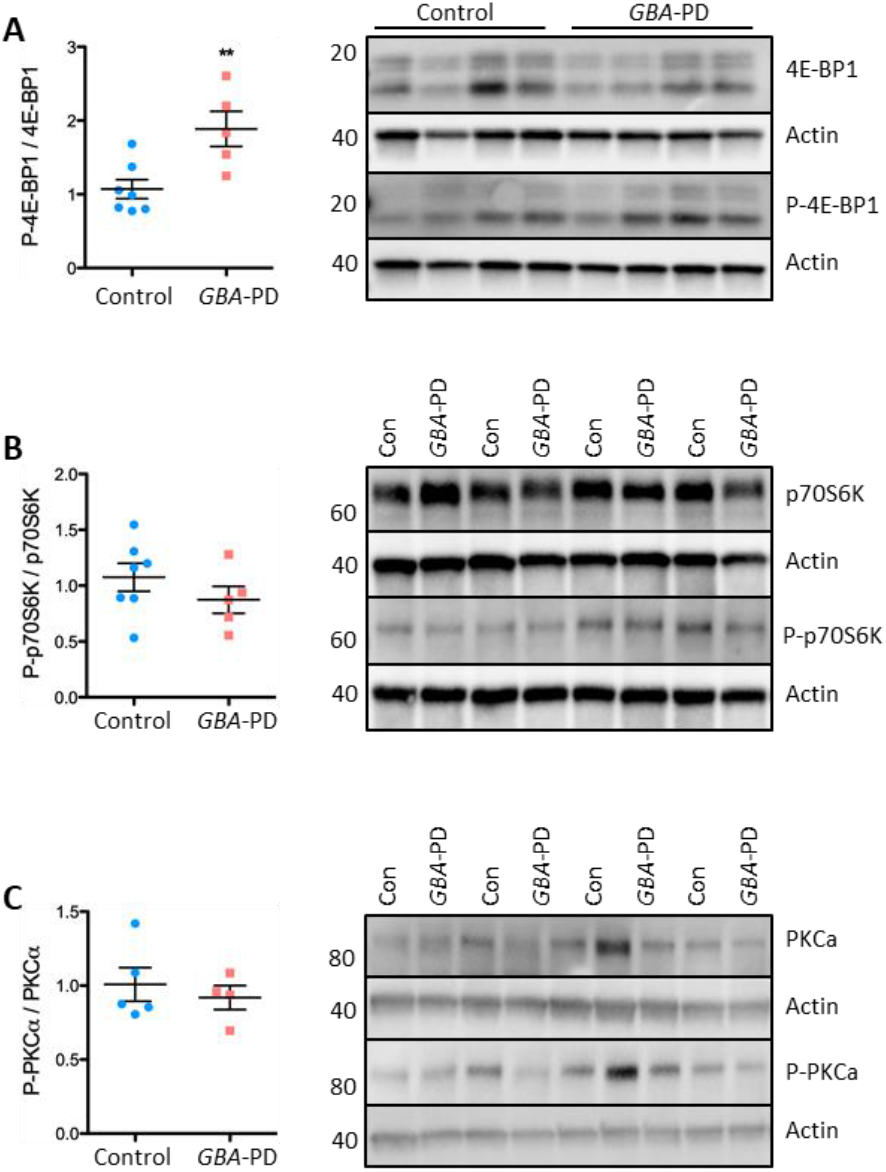
mTOR downstream effector 4E-BP1 is upregulated in *GBA*-PD patient neurons. A-C) Representative western blot and quantification of ratios of phosphorylated to total protein level of mTOR downstream targets (A) 4E-BP1 and phospho-4E-BP1 (P-4E-BP1), (B) p70S6K and phospho-p70S6K (P-p70S6K) and (C) PKCα and phospho-PKCα (P-PKCα) Expression levels normalized to β-actin and shown relative to control. Mean ± SEM, (n = 5-7 iPSC lines per group), **P≤0.01, (Student’s *t*-test).

**Fig. S6:**
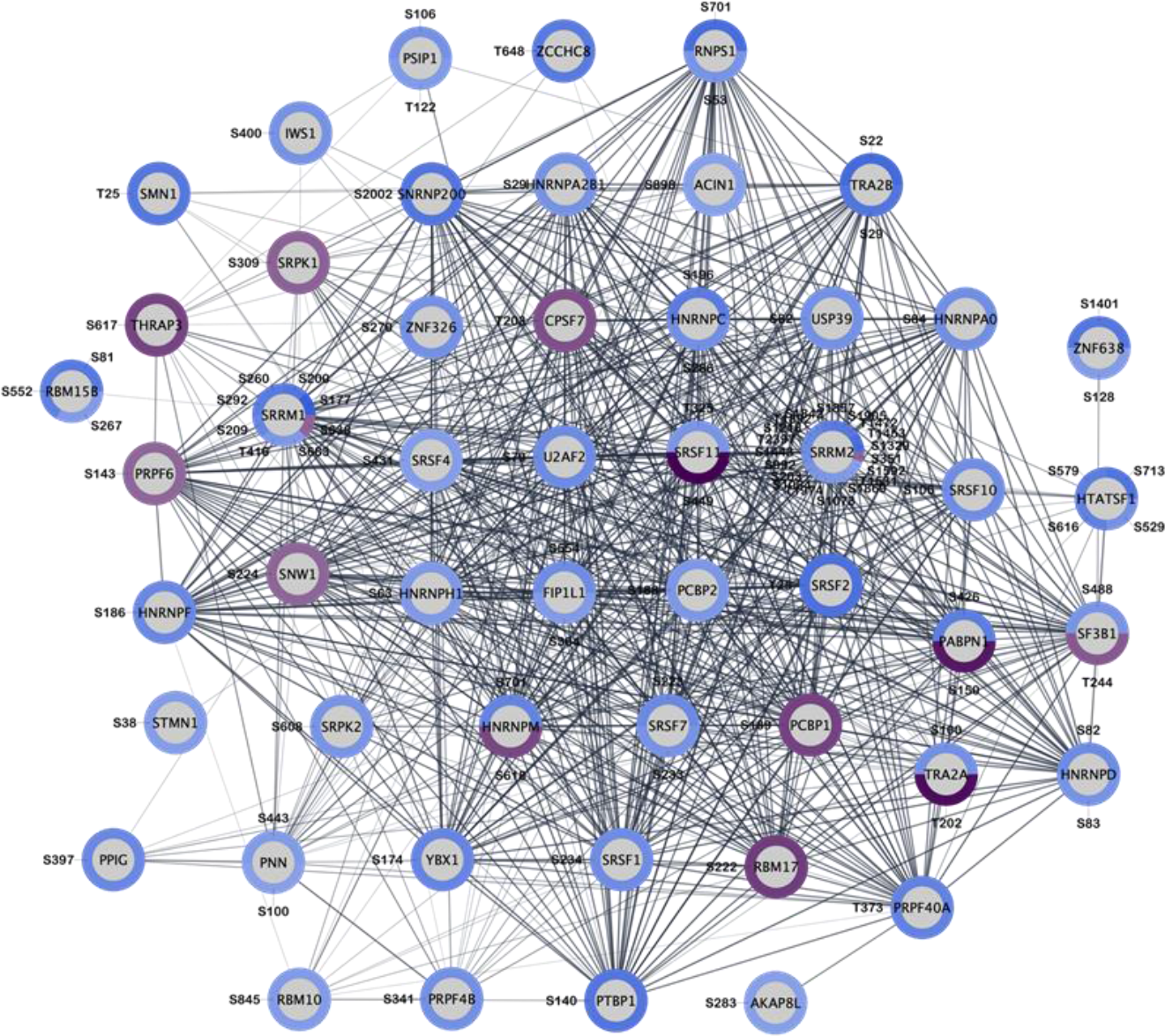
Proteins related to RNA splicing show altered phosphorylation states in *GBA* patient neurons. STRING network of functionally connected proteins related to the GO term “RNA splicing” with significantly regulated phospho-sites marked by the surrounding halo, where the colour signifies the phosphorylation abundance ratio (*GBA*-PD/control).

**Fig. S7:**
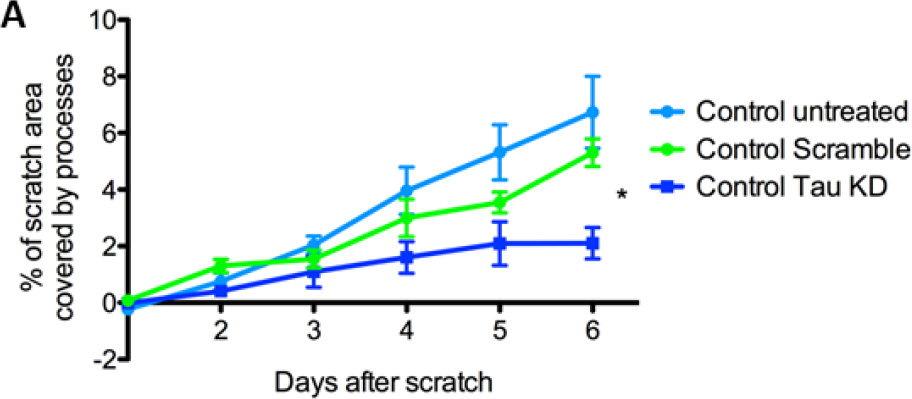
Knockdown of Tau in control neurons inhibits neurite outgrowth. A) Quantification of neurite outgrowth as measured by the percentage of the scratch area covered by processes. Data from control neurons (Con 3) with lentiviral-mediated shRNA tau KD, lentivirus with scrambled shRNA and untreated. Values from day 0 subtracted as background (n = 8-12 wells per group). Mean ± SEM, **P≤0.01, (One-way ANOVA, Dunnett’s multiple comparisons).

**Fig. S8:**
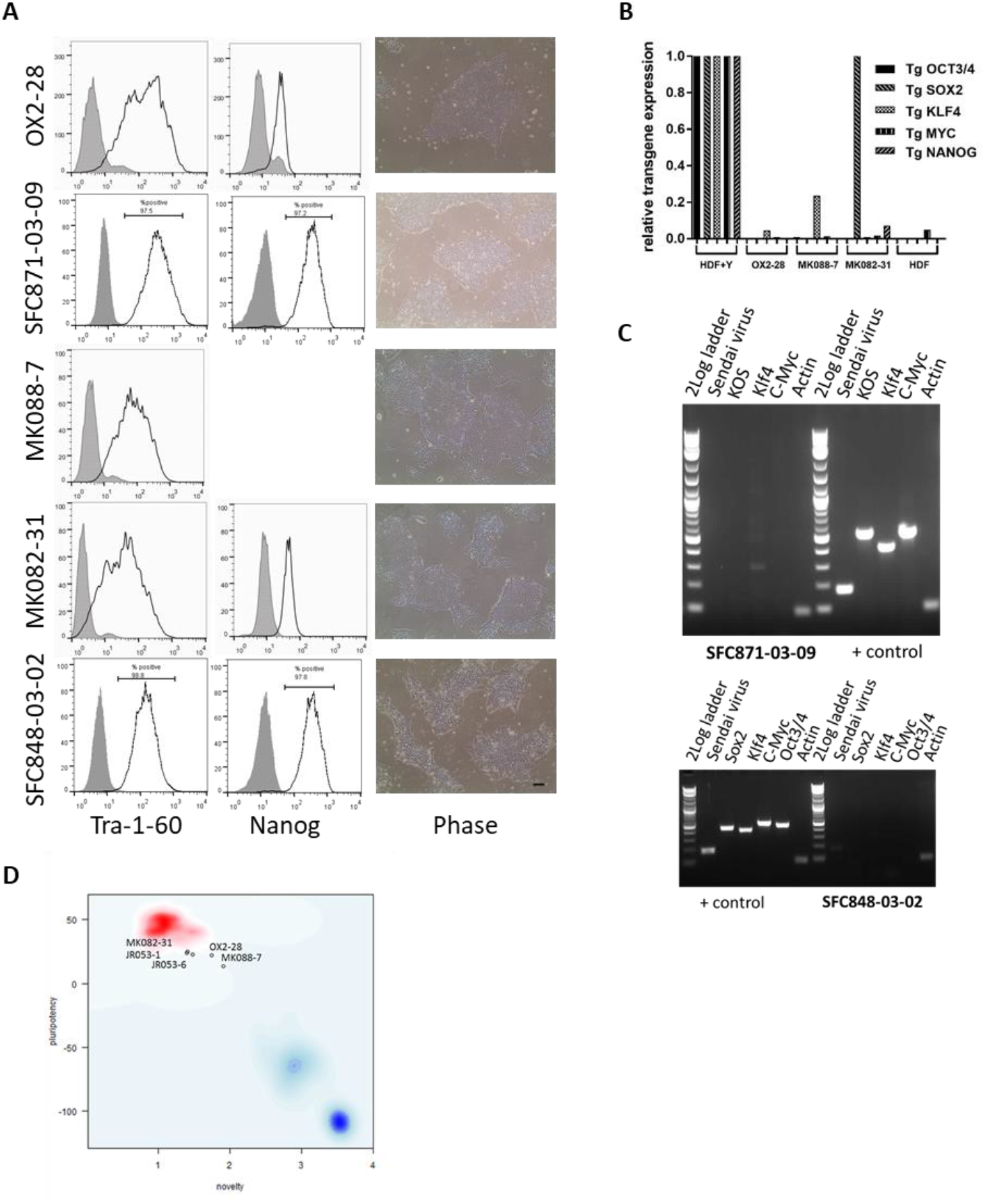
Reprogramming and characterization of iPSC lines from *GBA*-*N370S* patient and control fibroblasts. A) FACS analysis confirmed expression of pluripotency markers Tra-1-60 and Nanog in iPSCs; open black plot represents antibody, filled grey plot represents isotype control; Right-hand panel shows the expected iPSC colony morphology, with high nucleus to cytoplasm ratio by phase-contrast microscopy. Scale bar = 100 µm. B) Confirmation of retroviral transgene silencing with qRT-PCR analysis for each transgene, normalised to β-actin control, and expressed relative to levels in fibroblasts 5 days post-infection with the Yamanaka reprogramming retroviruses (HDF+Y). Uninfected fibroblasts (HDF) served as a negative control. C) Cytotune Sendai virus clearance in iPSC lines by qRT-PCR. L, Log2 ladder; Se, Sendai backbone 181 bp; S, Sox2 451 bp; K, Klf4 410 bp; M, c-myc 532 bp; O, Oct-4 483 bp; KOS, 528bp; A, β-actin control 92 bp; + positive control fibroblasts infected with Cytotune 5 days previously. iPSC lines show the correct size band for β-actin, and no bands corresponding to the reprogramming virus PCR product sizes. D) PluriTest analysis of Illumina HT12v4 transcriptome array data shows the tested iPSC lines cluster with the (previously published) control iPSC lines pluripotent stem cells in the red cloud and not with differentiated cells (blue clouds). Each circle represents one iPSC line.

**Fig. S9:**
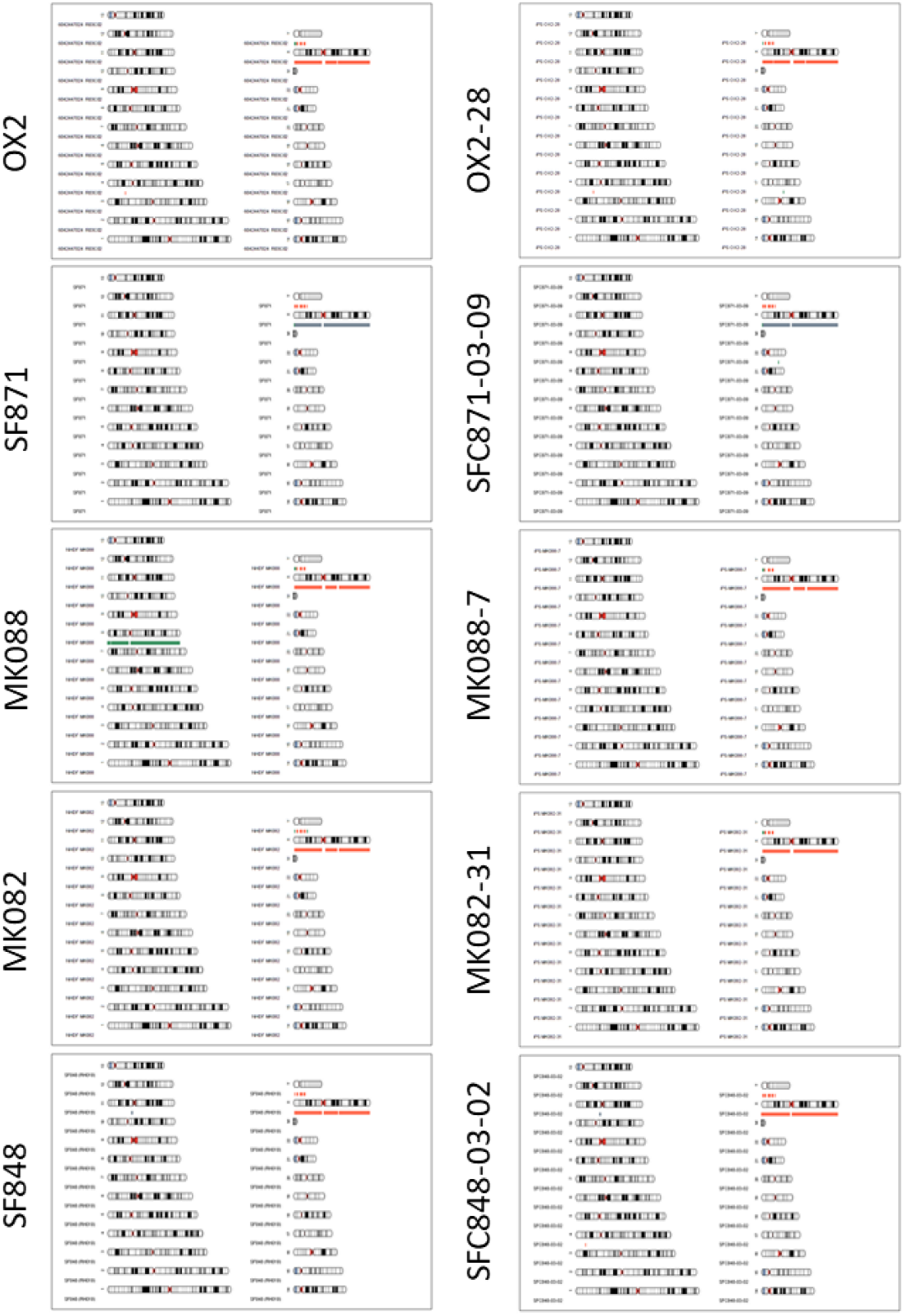
Karyotype analysis of unpublished iPSC lines used in the study. Genome integrity was assessed by Illumina Human CytoSNP-12v2.1 or OmniExpress24 SNP array and karyograms produced using KaryoStudio software (Illumina). Amplifications (green), deletions (orange) and LOH regions (grey) are shown alongside the relevant chromosome (except that in females the X chromosomes are annotated with grey, and single-copy sex chromosomes are annotated orange).

**Fig. S10:**
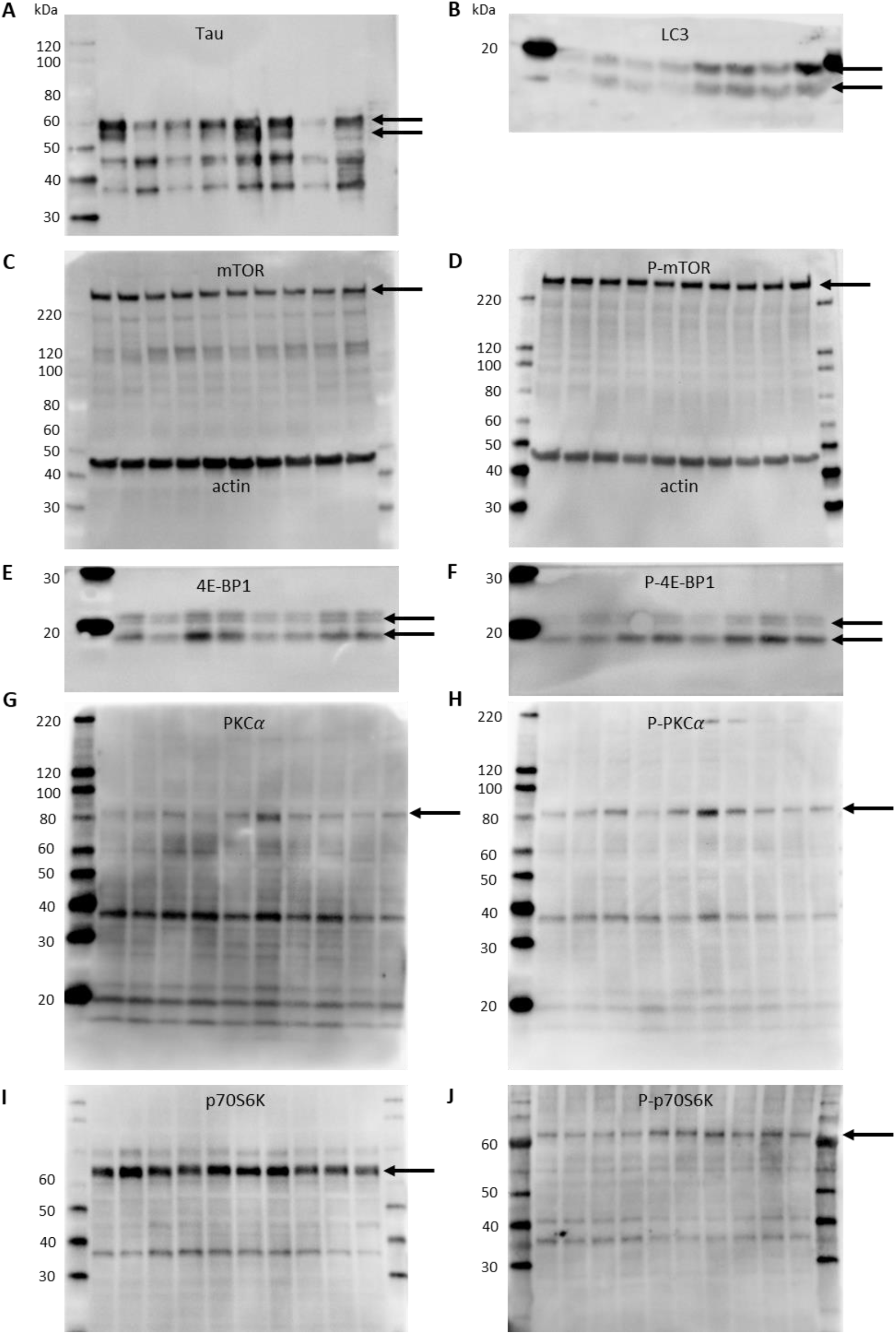
Representative Western blots for applied antibodies. A) microtubule-associated protein tau (Tau), B) microtubule-associated protein light chain 3 (LC3), C) mammalian target of rapamycin (mTOR), D) phospho-mTOR (P-mTOR), E) 4E-BP1, F) phospho-4E-BP1 (P-4E-BP1), G) PKCα and H) phospho-PKCα (P-PKCα), I) p70S6K, J) phospho-p70S6K (P-p70S6K). Arrows indicate the bands that were used for quantification.

**Table S1.** IPSC lines included in the study

| Study ID | Donor ID | Clone | Genotype | Age | Gender | Characterisation |
| --- | --- | --- | --- | --- | --- | --- |
| GBA 1 | MK088 | 07 | N370S/wt | 46 | M | This study |
| GBA 2 | SFC834-03 | 01 | N370S/wt | 72 | M | Fernandes et al. 2016 |
| GBA 3 | SFC871-03 | 09 | N370S/wt | 70 | F | This study |
| GBA 4 | MK082 | 31 | N370S/wt | 51 | M | This study |
| GBA 5 | MK071 | 03 | N370S/wt | 81 | F | Fernandes et al. 2016 |
| GBA 6 | SFC848-03 | 02 | N370S/wt | 68 | M | This study |
| Con 1 | SFC841-03 | 02 | wt/wt | 36 | M | Van Wilgenburg et al. 2013 |
| Con 2 | OX2 | 28 | wt/wt | 43 | M | This study |
| Con 3 | SFC856-03 | 04 | wt/wt | 78 | F | Haenseler et al. 2017 |
| Con 4 | MK053 | 06 | wt/wt | 68 | M | Lang et al. 2019 |
| Con 5 | OX1 | 19 | wt/wt | 36 | M | Van Wilgenburg et al. 2013 |
| Con 6 | NHDF | 1 | wt/wt | 44 | F | Hartfield et al. |

**Product code list:**
| Reagent | Company | Identifier |
| --- | --- | --- |
| **Cell culture** |  |  |
| hESC-Qualified Matrigel, LDEV-free | Corning | 354277 |
| 6-well plates | Corning | 3516 |
| mTeSR1 | Stem Cell Tech. | 5870 |
| Penicillin-streptomycin | ThermoFisher | 15070-063 |
| ROCK inhibitor | Bio-Techne | 1254 |
| TrypLE Express Enzyme | ThermoFisher | 12604 |
| Geltrex | ThermoFisher | A1413302 |
| Knockout DMEM | ThermoFisher | 10829018 |
| KnockOut serum replacement | ThermoFisher | 10828010 |
| β– mercaptoethanol | Sigma-Aldrich | M6250 |
| L-glutamine | ThermoFisher | 25030-032 |
| Non-essential amino acids | ThermoFisher | 11140-050 |
| Neurobasal Medium | ThermoFisher | 21103049 |
| N2 Supplement | ThermoFisher | 17502048 |
| B27 Supplement | ThermoFisher | 12587010 |
| Accutase | ThermoFisher | A11105-01 |
| Mitomycin C | Sigma-Aldrich | M4287 |
| LDN193189 | Sigma-Aldrich | SML0559 |
| SB431542 | Bio-Techne | 1614 |
| Sonic hedgehog-C24II | Bio-Techne | 1845-SH |
| Purmorphamine | Bio-Techne | 4551 |
| FGF8a | Strattech | 16124-HNAE-SIB |
| CHIR99021 | Bio-Techne | 4423 |
| BDNF | Peprtech | 450-02 |
| GDNF | Peprtech | 450-10 |
| TGFB3 | Peprtech | 100-36E |
| DAPT | Abcam | 120633 |
| Ascorbic acid | Sigma-Aldrich | A4544 |
| (db)-cAMP | Sigma-Aldrich | D0627 |
| **Proteomics** |  |  |
| Phosphate-buffered saline (PBS) | ThermoFisher | 14190 |
| Complete protease inhibitors | Roche | 11873580001 |
| PhosSTOP phosphatase inhibitors | Roche | 4906845001 |
| Urea | Sigma-Aldrich | U5128 |
| Thiourea | Sigma-Aldrich | 88810 |
| Sodium dodecyl sulfate | GE Healthcare | 17-1313-01 |
| **N-ethylmaleimide** | Sigma-Aldrich | E3876 |
| **Methanol** | Sigma-Aldrich | 34860 |
| **Chloroform** | Sigma-Aldrich | 288306 |
| **TCEP** | ThermoFisher | 77720 |
| **Endoproteinase Lys-C** | Wako | 129-02541 |
| **Triethylammonium bicarbonate** | Sigma-Aldrich | 17902 |
| **Trypsin** | Sigma-Aldrich | T0303 |
| **Trifluoroacetic acid** | Sigma-Aldrich | 106232 |
| **C18 3M Empore™ disk** | Sigma-Aldrich | 66883-U |
| **Poros 50 R2** | Applied Biosystems | 1-1159-06 |
| **Oligo R3** | Applied Biosystems | 1-1339-03 |
| **Acetonitrile** | Sigma-Aldrich | 271004 |
| **iTRAQ Reagent 8-plex assay kit** | Sigma-Aldrich | 4381662-1KT |
| **Glycolic acid** | Sigma-Aldrich | 124737 |
| **Titanium dioxide beads 10 µm** | GL Sciences | 5020-75010 |
| **Ammonium hydroxide solution** | Sigma-Aldrich | 17093 |
| **C8 3M Empore™ disk** | Sigma-Aldrich | 66882-U |
| **Dithiothreitol** | Sigma-Aldrich | D9163 |
| **N-glycosidase F** | Biolabs | P0705S |
| **Sialidase A** | Prozyme | GK80040 |
| **PhosSelect IMAC beads** | Sigma-Aldrich | P9740 |
| **Ammonium hydroxide** | Sigma-Aldrich | 17093 |
| **Immunofluorescence** |  |  |
| **96-well plates** | Greiner Bio-One | 655090 |
| **Paraformaldehyde** | Sigma-Aldrich | 158127 |
| **Distilled PBS (dPBS)** | ThermoFisher | 10010056 |
| **dPBS with calcium and magnesium** | ThermoFisher | 14040 |
| **Triton-X-100** | Sigma-Aldrich | T8787 |
| **Donkey serum** | Sigma-Aldrich | D9663 |
| **DAPI** | Sigma-Aldrich | D9542 |
| **rabbit anti-tyrosine hydroxylase** | Millipore | 152 |
| **goat anti-FOXA2** | R&D Systems | AF2400 |
| **mouse anti-β-III-Tubulin** | Biolegend | 801202 |
| **Western blotting** |  |  |
| **Tris-hydrochloride** | Sigma-Aldrich | T3253 |
| **Sodium chloride** | Sigma-Aldrich | S9625 |
| **Tergitol-type NP-40** | Sigma-Aldrich | NP40S |
| **Sodium cholate hydrate** | Sigma-Aldrich | C9282 |
| **Sodium dodecyl sulfate** | Sigma-Aldrich | BCBN2052V |
| **Bicinchoninic acid assay** | BCA, Pierce | 23225 |
| **4-15% precast gel** | Biorad | 567-8085 |
| **Trizma base** | Sigma-Aldrich | T1503 |
| **Polyvinylidene difluoride membrane** | Biorad | 170-4157 |
| **Glycine** | Sigma-Aldrich | G8898 |
| **MagicMark XP Protein Standard** | ThermoFisher | LC5602 |
| **BLUeye Prestained Protein Ladder** | Geneflow | S6-0024 |
| **Skim milk** | Sigma-Aldrich | 70166 |
| **Tween 20** | Sigma-Aldrich | P2287 |
| **Bovine-serum albumin** | Sigma-Aldrich | A9418 |
| **HRP-conjugated secondary antibody** | BioRad | 170-6515 |
| **ECL** | Millipore | P90720 |
| **Mouse anti-b-Actin-HRP** | Abcam | 49900 |
| **rabbit anti-LC3** | Sigma-Aldrich | L7543 |
| **mouse anti-Tau** | Neomarkers | ms-247-P |
| **mouse anti-mTOR** | Cell Signaling | 45175 |
| **rabbit anti-phospho-mTOR Ser2448** | Cell Signaling | 2971 |
| **rabbit anti-p70S6K** | Cell Signaling | 2708 |
| **mouse anti-phospho-p70S6K Thr389** | Cell Signaling | 9206 |
| **rabbit anti-4E-BP1** | Cell Signaling | 4923 |
| **rabbit anti-phospho-4E-BP1 Thr37/46** | Cell Signaling | 2855 |
| **rabbit anti-PKCα** | Cell Signaling | 2056 |
| **rabbit anti-phospho-PKCa Ser657** | ThermoFisher | PA5-78124 |
| **Lenti-virus production** |  |  |
| **Gibson Assembly Master Mix** | NEB | E2611 |
| **KAPA HiFi HotStart ReadyMix** | KAPA Biosystems | KK2602 |
| **QIAquickGel Extraction Kit** | QIAGEN | 28704 |
| **One Shot TOP10 Chemically Competent E. coli** | ThermoFisher | C40400 |
| **One Shot Stbl3 Chemically Competent E. coli** | ThermoFisher | C737303 |
| **PureLink HiPure Plasmid Maxiprep kit** | ThermoFisher | K210006 |
| **Calcium chloride** | Sigma-Aldrich | C4901 |
| **HEPES** | Sigma-Aldrich | H3375 |
| **Sodium phosphate dibasic** | Sigma-Aldrich | 255793 |
| **Sodium butyrate** | Sigma-Aldrich | B5887 |
| **Sucrose** | ThermoFisher | S/8600/60 |
| **D-mannitol** | Sigma-Aldrich | M9546 |
| **Assays** |  |  |
| **Citric acid** | Sigma-Aldrich | C2404 |
| **Dibasic sodium phosphate** | Sigma-Aldrich | S7907 |
| **Taurocholic acid** | Sigma-Aldrich | T4009 |
| **Conduritol B epoxide** | Calbiochem | 234599 |
| **Methylumbellifery β-D-glucopyranoside** | Sigma-Aldrich | M3633 |
| **NCGC00188758** | Sigma-Aldrich | 5316600001 |

## Notes

### Competing Interest Statement

The authors have declared no competing interest.

